# *Trypanosoma cruzi* isolates naturally adapted to congenital transmission display a unique strategy of transplacental passage

**DOI:** 10.1101/2022.06.30.498325

**Authors:** Paula Faral-Tello, Gonzalo Greif, Selva Romero, Andrés Cabrera, Cristina Oviedo, Telma González, Gabriela Libisch, Ana Paula Arévalo, Belén Varela, José Manuel Verdes, Martina Crispo, Yester Basmadjián, Carlos Robello

## Abstract

Chagas disease is mainly transmitted by vertical transmission (VT) in non-endemic areas and in endemic areas where vector control programs have been successful. For the present study, we isolated natural strains vertically transmitted through three generations and proceeded to study their molecular mechanism of VT. No parasitemia was detected in immunocompetent mice, but they were able to induce an immune response and colonize different organs. VT experiments revealed that infection with the different strains did not affect mating, pregnancy or resorptions, but despiting low parasitemia, VT strains reached the placenta and resulted in higher vertical transmission rates than strains of either moderate or high virulence. While the virulent strain modulated more than 2500 placental genes, VT strains modulated 150, and none of the modulated genes is shared between them. VT strains downregulate genes associated with cell division and replication and upregulate immunomodulatory genes leading to anti-inflammatory responses and tolerance. The virulent strain stimulates a strong pro-inflammatory immune response, and this molecular footprint correlated with histopathological analyses. We herein describe a unique placental response regarding the passage of T cruzi VT isolates across the maternal-fetal interphase, challenging the current knowledge derived mainly from studies of laboratory-adapted or highly virulent strains.

## 1. Introduction

The acronym TORCH (toxoplasmosis, rubella, cytomegalovirus, and herpes simplex virus) refers to those pathogens that are transmitted in the human species from mother to offspring through transplacental passage. The referred acronym is more exclusive than inclusive, leaving out an expanding list of pathogens that are vertically transmitted and represent severe human health problems. In recent years, the relevance of this transmission mechanism has become clear, and Chagas disease (CD) is a paradigmatic case. In the absence of the vector, *Trypanosoma cruzi* can still be transmitted through blood transfusions, organ transplant or congenitally. Hence the migration of asymptomatic individuals from endemic to non-endemic areas due to poverty conditions in the context of globalization has led to its current consideration as a re-emerging disease (Guarner, 2019). Unlike vectorial transmission, all other forms of transmission are possible in any country (Gascon et al., 2010), and the World Health Organization estimates that at least one-quarter of the total burden of newly reported cases corresponds to Congenital Chagas Disease (CCD) (WHO, 2015), which includes endemic areas with successful vectorial control programs and non-endemic areas where vertical transmission (VT) is the only active route of transmission. Although CD is considered a vector-borne disease, these events have radically changed its epidemiological profile, turning it into a global urban disease and a worldwide health problem (Oliveira et al., 2010; Lidani et al., 2019). The rates of *T. cruzi* vertical transmission range from 1% to 12%, depending on the area (Bustos et al., 2019), with an average rate of 5% (Howard et al., 2014). CCD often presents as an acute infection and although 60% of infected babies are born asymptomatic, they usually exhibit detectable levels of parasitemia and display higher frequencies of low weight, prematurity, and lower APGAR scores (Aspect, Pulse, Grimace, Activity and Respiration) when compared to non-infected babies (Carlier and Truyens, 2015; Amorín and Pérez, 2016; Antinori and Corbellino, 2018). Although rarely lethal, untreated CCD can lead to hepatosplenomegaly, meningoencephalitis, and myocarditis (Bittencourt et al., 1975); however, when detected and treated early, parasitic clearance is more than 90% effective (Schijman et al., 2003; Altcheh et al., 2005). One important aspect of CCD is that most infected pregnant women are diagnosed for CD during routine controls, because they are usually asymptomatic (Howard et al., 2014). This observation led us to the hypothesis that there could be natural strains adapted to vertical transmission, capable of persisting in a ‘silent’ state in the host and that are able to cross the placental barrier during pregnancy. In addition, these microorganisms would constitute a successful case of parasitism and are not accurately modeled by highly virulent and laboratory adapted strains. To test the hypothesis, we identified familial cases of Congenital Chagas in which *T. cruzi* strains have asymptomatically passed through generations: from great-grandmothers, to grandmothers, then to mothers and their children. By xenodiagnoses of babies born to positive mothers, we isolated and investigated the strains in terms of virulence, tissue tropism, as well as through the development of a vertical transmission murine model coupled to transcriptomic placental response.

## 2. Materials and methods

### 2.1. Clinical cases and diagnosis

*<u>Case 1</u>* (2013): asymptomatic 12-month-old baby born and living in a non-endemic area (Montevideo, UY) with a positive mother (24 years old) who also lived in a non-endemic area. Positive grandmother and great grandmother living in a non-endemic area (Montevideo) and in an endemic area (Salto, UY), respectively; weight and gestational age at birth: 3.400 kg and 38 weeks. *<u>Case 2</u>* (2015): asymptomatic 1-month-old baby born in a non-endemic area (Montevideo, UY) to a positive mother, also born and living in Montevideo, who had two other positive children; weight and gestational age at birth: 3.910 kg and 39 weeks. *<u>Case 3</u>* (2017): a 3-month-old baby born asymptomatic in a non-endemic area, who developed an atrioventricular blockage at 3 months of age; positive mother; weight and gestational age at birth: 3.320 kg and 41 weeks.

All mothers and babies were born in non-endemic areas. Xenodiagnosis (5 inbred stage IV nymphs of *Triatoma infestans*) was performed in babies, and after bites, nymphs’ feces were monitored at 20, 40 and 60 days post diagnosis. The procedure was repeated every 6 months until the baby was either positive or 18 months old. Xenodiagnosis was considered positive when *T. cruzi* compatible trypomastigotes were observed in at least one nymph. Congenital infection was inferred due to patients residing in non-endemic areas and never having traveled or received blood transfusions. A summary of the characteristics of each clinical case is given in Table 1.

**Table 1.**
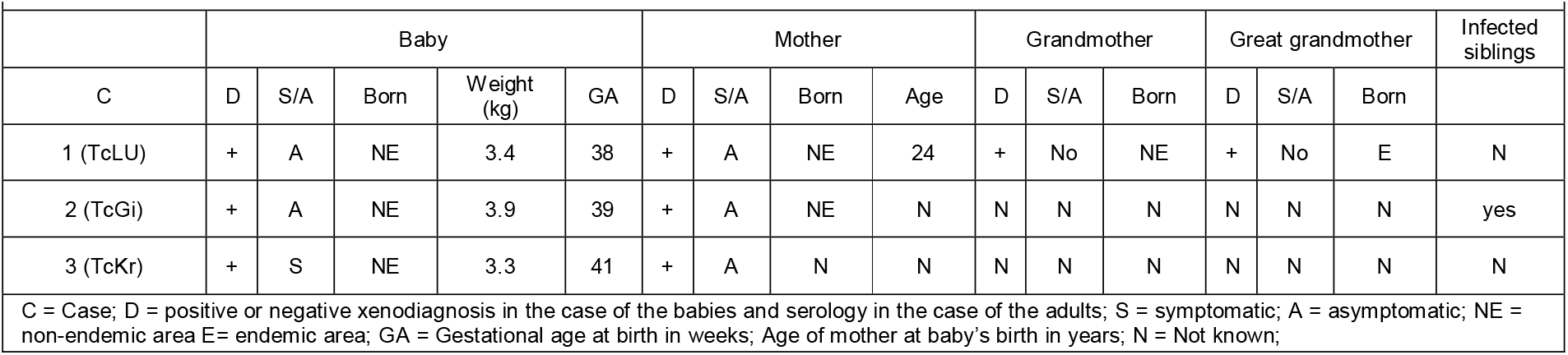
Summary of characteristics of the clinical cases from which VT strains were isolated

|  |  |  |  |  |  |  |  |  |  |  |  |  |  |  |  |  |
| --- | --- | --- | --- | --- | --- | --- | --- | --- | --- | --- | --- | --- | --- | --- | --- | --- |
|  | Baby |  |  |  |  | Mother |  |  |  | Grandmother |  |  | Great grandmother |  |  | Infected siblings |
| C | D | S/A | Born | Weight (kg) | GA | D | S/A | Born | Age | D | S/A | Born | D | S/A | Born |  |
| 1 (TcLU) | + | A | NE | 3.4 | 38 | + | A | NE | 24 | + | No | NE | + | No | E | N |
| 2 (TcGi) | + | A | NE | 3.9 | 39 | + | A | NE | N | N | N | N | N | N | N | yes |
| 3 (TcKr) | + | S | NE | 3.3 | 41 | + | A | N | N | N | N | N | N | N | N | N |

C = Case; D = positive or negative xenodiagnosis in the case of the babies and serology in the case of the adults; S = symptomatic; A = asymptomatic; NE = non-endemic area E= endemic area; GA = Gestational age at birth in weeks; Age of mother at baby's birth in years; N = Not known;

### 2.2. Animals

Inbreed BALB/cJ and Nu/J 8-week-old females (Jackson Laboratory, Bar Harbor, US) were housed under specific pathogen-free conditions in individually ventilated cages (IVC, Tecniplast, Milan, Italy) at the Animal Facility of the Unit of Biotechnology of Laboratory Animals. During infection, mice were assigned to groups of no more than six and housed in a Biosafety Level 2 (BSL2) ventilated rack with HEPA filters (IsoCage N, Tecniplast). All infection procedures were performed under a BSL2 laminar flow hood to comply with biosafety institutional rules. Animals received autoclaved fresh water and a standard autoclaved mouse diet *ad libitum* (Labdiet 5K67, PMI Nutrition, IN, US).

### 2.3. Strain isolation and maintenance

Feces of positive nymphs were resuspended in salinated water and inoculated into BALB/cJ mice (Jax strain # 000651) via intraperitoneal route (ip). Parasitemia was assessed by microscopic observation of blood samples obtained by submandibular puncture every 3 days. When parasitemia was detected, an immunosuppression protocol with cyclophosphamide (Filaxis) was applied at 40 mg/kg/day for 5 days via ip route. Afterwards, complete blood was used to inoculate three Nu/J mice (Jax strain # 002019) to amplify the parasite load and create parasite stocks for further use in the experiments. Nu/J mice were infected through ip route with 2×10^5^ parasites, and parasitemia was monitored 2-3 times a week by blood parasite counting obtained by puncture of the submandibular vein until a parasitemia peak of 5×10^6^ parasites/mL was reached. Thereafter, mice were deeply anesthetized via ip with a mixture of ketamine 100mg/kg (Vetanarcol®, König, Buenos Aires, Argentina) and Xylazine 10mg/kg (Seton®, Calier, Barcelona, Spain) for final bleeding. Blood was collected by cardiac puncture and mice were immediately euthanized by cervical dislocation. Total parasites were purified by differential centrifugation in a Percoll gradient as described elsewhere (Grab and Bwayo, 1982) and counted to be used in further experiments. The first parasitemia peak in nude mice was considered passage 1 (P1), and experiments were always performed with P1 to P5 parasites to conserve biological features. P1 to P5 *T. cruzi* VT isolates from Case 1 (TcLu), Case 2 (TcGi) and Case 3 (TcKr) and trypomastigotes from the Dm28c (DTU TcI) and Garbani (DTU TcVI) strains were purified every time from nude mice blood. Fresh parasites were used in all the experiments conducted in this study.

### 2.4. Molecular typing

Purified DNA from all strains was obtained from trypomastigotes and was used to identify the Discrete Typing Unit (DTU) based on specific PCR products: the intergenic region of spliced leader genes (SL-IR), the 24Sα ribosomal DNA subunit (rDNA 24Sα), and the A10 fragment previously described (Burgos et al., 2007). PCR products were seeded in 5% MetaPhor® agarose gels, and amplification products were compared to reference strains. PCR products were purified and verified by Sanger sequencing. DTU determination was afterwards confirmed by Dr. Alejandro Shijman (Instituto de Investigaciones en Ingeniería Genética y Biología Molecular ‘Dr. Héctor N. Torres’ INGEBI-CONICET, Buenos Aires, Argentina) by a multiplex RealTime PCR developed by their group (Cura et al., 2015).

### 2.5. Epimastigogenesis

Trypomastigotes purified from mice blood were incubated in PBS buffer (pH 6) at 37°C for four hours before epimastigogenesis in Liver Infusion Tryptoese (LIT) as previously described (Kessler et al., 2017). Different time points were collected to count epimastigote forms and for immunofluorescence staining.

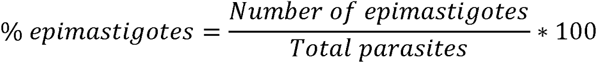

### 2.6. Cell culture

HFF-1 (Human Foreskin Fibroblasts, SCRC-1041™) were purchased from ATCC®, and SWAN-71 cells (Straszewski-Chavez et al., 2009) were kindly donated by Dr. Gil Mor (C.S Mott Center for Human Growth and Development, Department of Obstetrics and Gynecology, Wayne State University). Both cell lines were grown in Dulbecco’s Modified Eagle medium (DMEM) (Life Technologies, Grand Island, NY, US) supplemented with 10% FBS at 37°C in a 5% CO2 humidified atmosphere. The epimastigote forms used were grown in LIT supplemented with 10% FBS at 28°C.

### 2.7. Determination of infectivity index in mammalian cells

Nude mice trypomastigotes derived from all strains were purified, counted and incubated with HFF-1 and SWAN-71 semi-confluent monolayers for 2 hours at a multiplicity of infection of 5 parasites to 1 host cell. Afterwards, non-internalized trypomastigotes were washed 5 times with PBS and cells were incubated for another 48 hours before immunofluorescence microscopy. After 48 hours, coverslips were washed with PBS, fixed with 95% (v/v) ethanol and stained with Fluoroshield™ with DAPI (Sigma, US). Infectivity was evaluated considering invasion and replication capacity, counting infected cells and parasites per infected cell in the infection photos obtained from each experimental condition. For each replicate, a total of at least 500 cells were counted and the results were expressed in graphs as the mean and SE of 3 independent experiments. The infection index was calculated as:

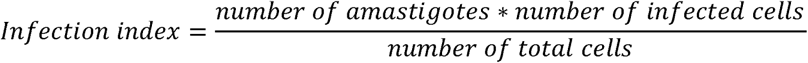

### 2.8. Indirect immunofluorescence

Infected cells plated on coverslips or parasite pellets were washed with PBS and fixed with PBS-4% paraformaldehyde, permeabilized for 5 minutes with 0.1% Triton X-100 and blocked for 30 minutes with 5% bovine serum albumin (BSA) (all reagents purchased in Sigma). Cells from infectivity and replications assays were incubated 1 hour at room temperature with anti-cytosolic tryparedoxin peroxidase (cTXNPx) primary antibody (Pineyro et al., 2008) and 30 minutes with Alexa Fluor 488 anti-rabbit IgG secondary antibody (Invitrogen, Eugene OR). Parasites from the epimastigogenesis experiment were attached to poly-L-lysine-coated slides and treated as monolayers, except for those incubated with monoclonal primary antibody (Mab25, kindly provided by Dr. Sergio Schenkman, 1:1000 dilution) that binds to the *T. cruzi* flagellar calcium-binding protein (Schenkman et al., 1991). All antibodies were diluted in 1% BSA/PBS and coverslips and parasites mounted with ProLong Antifade reagent.

### 2.9. Mice infection, monitoring, survival and parasitemia curves

For mice survival and parasitemia curves, groups of six 6-8-week-old BALB/cJ females were inoculated with Nu/J derived trypomastigotes and parasitemia was monitored as described above. An uninfected control group was included in every experiment. Initial trypomastigote dosages were 5×10^4^ and 1×10^4^ for the survival and parasitemia curves except for Garbani in which case dosages were 10^4^ and 10^3^ due to its virulence. On post-infection (pi) day 35, when parasites were not visually detected in blood, mice were anesthetized by intraperitoneal inoculation of a mixture of ketamine 100mg/kg and Xylazine 10mg/kg. Blood was collected by cardiac puncture, and mice were euthanized by cervical dislocation. Blood was placed into dry or heparinized tubes for serum collection or molecular testing, respectively. Organs (heart, gut, spleen and uterus) were perfused with ice-cold PBS, and except for a piece of the spleen that was dissected and immediately stored at -80°C in TRIzol Reagent (Invitrogen) until RNA extraction, the rest of the spleen and organs were dry-frozen at -80°C for further DNA extraction.

### 2.10. Evaluation of mating, pregnancy, implantation rate and vertical transmission

For vertical transmission experiments, 8-week-old BALB/cJ female mice were randomly separated into six groups (six mice per group) and infected with the different strains, an uninfected control group was also included. Initial inoculums were 5×10^4^ for VT strains, 1×10^4^ for Dm28c, and 1×10^3^ for Garbani. Injection and monitoring were performed as described above. To evaluate the reproductive parameters and vertical transmission, 30 days after infection, infected females were placed for 2 days in cages with dirty bedding from males (Whitten effect) to synchronize their estrous cycles. Afterwards, mating was performed by adding one proven male in a cage with 3 females for 5 days. Females were checked every day for the presence of a vaginal plug (VP), and the average of VP positive females registered during those 5 days was used to calculate mating percentage as described next. All females were anesthetized and euthanized on day 20 of gestation, and viable fetuses and resorptions were recorded to measure pregnancy and implantation rates. Maternal blood, heart, gut, spleen, uterus, and fetal, placental and resorption tissues were collected and processed according to the experiment procedure for each case.

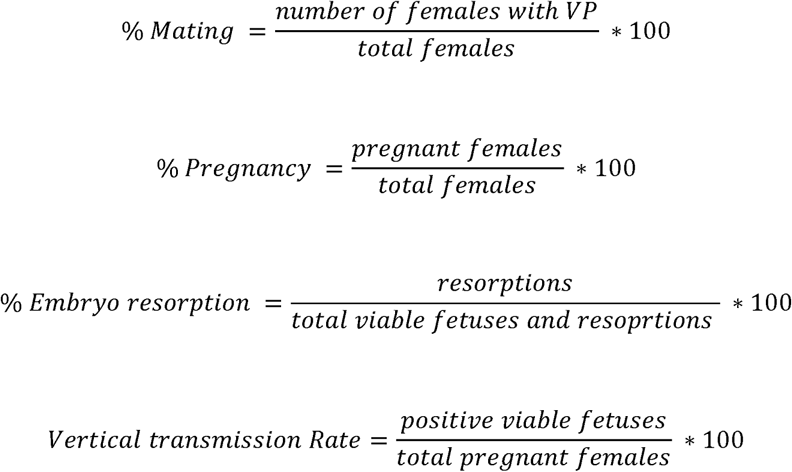

### 2.11. *T. cruzi* DNA detection by qPCR

Blood DNA was extracted using *Quick*-DNA™ Miniprep Plus Kit (D4069, Zymo Research) and organ DNA, using *Quick*-DNA™ Tissue/Insect Plus Kit (D6016, Zymo Research) according to manufacturer’s recommendations. Tissue disruption was achieved using the Bullet Blender® Tissue homogenizer (Next Advanced) and the kit of beads. Total DNA was quantified, diluted to 100ng/µL, and 1 µL of the sample was amplified using a kDNA TaqMan probe (Ramirez et al., 2015) with an IDT mix in a final volume of 10 µL, and the relative expression was calculated by delta-delta Ct method and normalized against the mouse *gapdh* housekeeping gene. Samples were analyzed in duplicate using the QuantStudio™ 3 System thermocycler (Applied Biosystems-Thermo Fisher Scientific). Primers and probe sequences are detailed in Supplementary Table 1. On the plate, reaction controls with water, positive controls with *T. cruzi* gDNA and negative control with DNA from uninfected mice were included. The latter was used to set up a negative cut-off line for PCR results in infected organs.

### 2.12. Cytokine expression

Mice infected with *T. cruzi* were euthanized at 30 and 60 dpi, and the spleens were removed and subjected to RNA extraction with TRI Reagent (Invitrogen) and Direct-zol RNA Miniprep Plus kit (R2072, Zymo Research) according to manufacturer’s instructions. cDNA was synthesized from 500ng of RNA using SuperScript II retrotranscriptase (Invitrogen) using Oligo(dT). Reactions were performed in a final volume of 10 μL using SybrGreen (KAPA SYBR Fast Universal 2x qRT-PCR master mix), a primer final concentration of 200nM and 1μL of a 1/5 dilution of the cDNA. Each reaction was performed in duplicate in QuantStudio 3 System thermocycler (Applied Biosystems, Thermo Fisher Scientific). Melt curves were generated to check specific amplification products, and Ct values obtained for each gene were normalized against the expression of the *Mus musculus gapdh* gene. Delta Ct was determined to calculate the relative expression for each gene in every sample by the delta-delta Ct method. Primers sequences used to amplify the different cytokines are listed in Supplementary Table 1.

### 2.13. Transcriptomics

For the RNAseq experiment, three placentas for each condition (biological replicates) were selected and processed to extract total RNA. Briefly, -80°C stored placentas were disrupted in Tri-Reagent (Thermo) in a Bullet Blender (Next Advance) with 1.6mm stainless steel beads (SSB16, Next Advance) for 10 minutes at maximum velocity. Once the tissue was disrupted, 200 µL of chloroform for 1 mL of Tri-Reagent was added, homogenized and centrifuged for 15 minutes at 15000rcf. The aqueous phase was recovered and processed with a Direct-zol RNA kit (Zymo Research, USA) according to the manufacturer’s instructions. RNA quality was determined using the Bioanalyzer RNA 6000 Nano kit (Agilent, USA) in a Bioanalyzer 2100 (Agilent, USA). An RNA Integrity Number (RIN) higher than 8 was considered acceptable to continue to library construction and sequencing. Library construction and sequencing were performed in Macrogen (Korea). For library construction, TruSeq stranded total RNA with Ribo-Zero Gold (Illumina, USA) was used. Sequencing was performed in the Illumina NovaSeq6000 platform. Paired-end reads of 100 base pairs were obtained. More than 60 million reads were obtained for each sample (Supplementary Table 2). Raw data were deposited in NCBI (SRA) under accession number PRJNA820598. Sequences obtained were quality checked using FastQC (Andrews, 2010). Reads were mapped against Mouse Genome version 39 (GRCm39.primary_assembly genome download from Ensembl) with a STAR package (Dobin et al., 2013) using annotation file M26 released by the Gencode project. Reads counts were performed using FeatureCounts (Liao et al., 2014), mapping statistics and STAR and FeatureCounts parameters were included in Supplementary Table 2. Feature counts output was processed in RStudio using DESeq2 package (Love et al., 2014) to obtain differential expression genes (DEG) using |log2FC| > 1.5 and Benjamini-Hochberg adjusted p-value < 0.05 (or <0.01 when indicated) to perform downstream analysis. A table of differentially expressed genes (DEG) was generated for each condition as compared to the control group (uninfected animals) (Supplementary Table 3). DEG clusterization was performed using pheatmap package in RStudio software or GraphPrism v9.0. Gene Ontology analyses were performed in Metascape (Zhou et al., 2019).

### 2.14. Histopathology

Immediately after the autopsy, whole placentas were fixed in 10% neutral buffered formalin (pH 7.4) for further processing. After fixation, the placentas were transversally hemi-sected, embedded in paraffin, sectioned in 5 μm slices and stained with hematoxylin–eosin (H&E) according to Sala et al. (2012). For histological evaluation, hemisected placentas were examined under a light microscope (BX41, Olympus) at 10× magnitude, considering the selected regions: maternal decidua basalis, trophoblastic giant cells zone, spongiotrophoblasts zone and labyrinth zone, according to Georgiades et al. (2002). Each specimen was evaluated by two different pathologists (BV, JMV) to establish a histopathological score in each case.

### 2.15. Ethical statement

Experimental animal protocols containing all procedures reported here were opportunely approved by the Institutional Animal Ethics Committee (protocol numbers CEUA 007-17 and CEUA 010-17), and in accordance with National Law 18.611 and the International Guide for the Care and Use of Laboratory Animals (National Research Council (US) Committee, 2011).

## 3. Results

### 3.1. Molecular typing of isolates

Xenodiagnostic tests were performed on three newborns whose mothers and grandmothers were seropositive for CD, and after 15-25 days, parasites were observed in insect feces. Insect intestinal homogenates were inoculated into BALB/cJ mice, and parasites in blood were detected after 30-45 days, although in all cases they were observed after counts of at least one hundred microscopic fields. Parasitemia peaks were obtained only after immunosuppression, and three isolates were obtained and named TcLU, TcKR and TcGI. Thereafter, due to their very low virulence in immunocompetent mice, all of the isolates were maintained by passages on Nu/J mice which did not control the infection. Molecular typing indicated that TcGI, TcKR and TcLU belonged to the TcV DTU or BCb lineage (Berna et al., 2021) (**Supplementary figure 1**).

### 3.2. Murine infection assays indicate that VT isolates reproduce their clinical features in the murine model

To compare the infection course and virulence of the different strains, Kaplan-Meier and parasitemia curves with different parasite loads were performed on BALB/cJ mice. As shown in the survival curves (**Figure 1A**), 100 per cent survival was observed for both the Dm28c and VT strains when 5×10^4^ trypomastigotes were inoculated. On the other hand, when inoculated with 1×10^4^ parasites of the Garbani strain, 100% of the mouse population reached the ethics endpoint approximately by day 19 pi, and with an inoculum of 1×10^3^ trypomastigotes, 50% of the mice reached the endpoint by day 29. As shown in **Figure 1B**, an initial inoculum of 5×10^4^ parasites of the Dm28c strain generates a peak of blood parasitemia of 4×10^5^ parasites per mL at day 20 pi, and by the 36th day, no parasites were observed. Initial inoculums of 1×10^4^ with the Garbani strain exhibited a peak of parasitemia of 7×10^5^ parasites/mL on day 12 pi from which the animals did not recover (**Figure 1A**). When the Garbani inoculum was reduced to 1×10^3^, 50% of the surviving mice showed a parasitemia curve with a peak of 4×10^5^ parasites per mL on day 10 pi, and parasite clearance from blood was observed on day However, for the VT strains, parasitemia was almost undetectable (**Figure 1B**) when 5×10^4^ parasites were inoculated. These results led us to the conclusion that the VT strains present low virulence and a silent phenotype in mice, whereas the Garbani and Dm28c clones exhibit high and medium virulence, respectively, which is the reason why they were used for comparative studies throughout this work.

**Figure 1.**
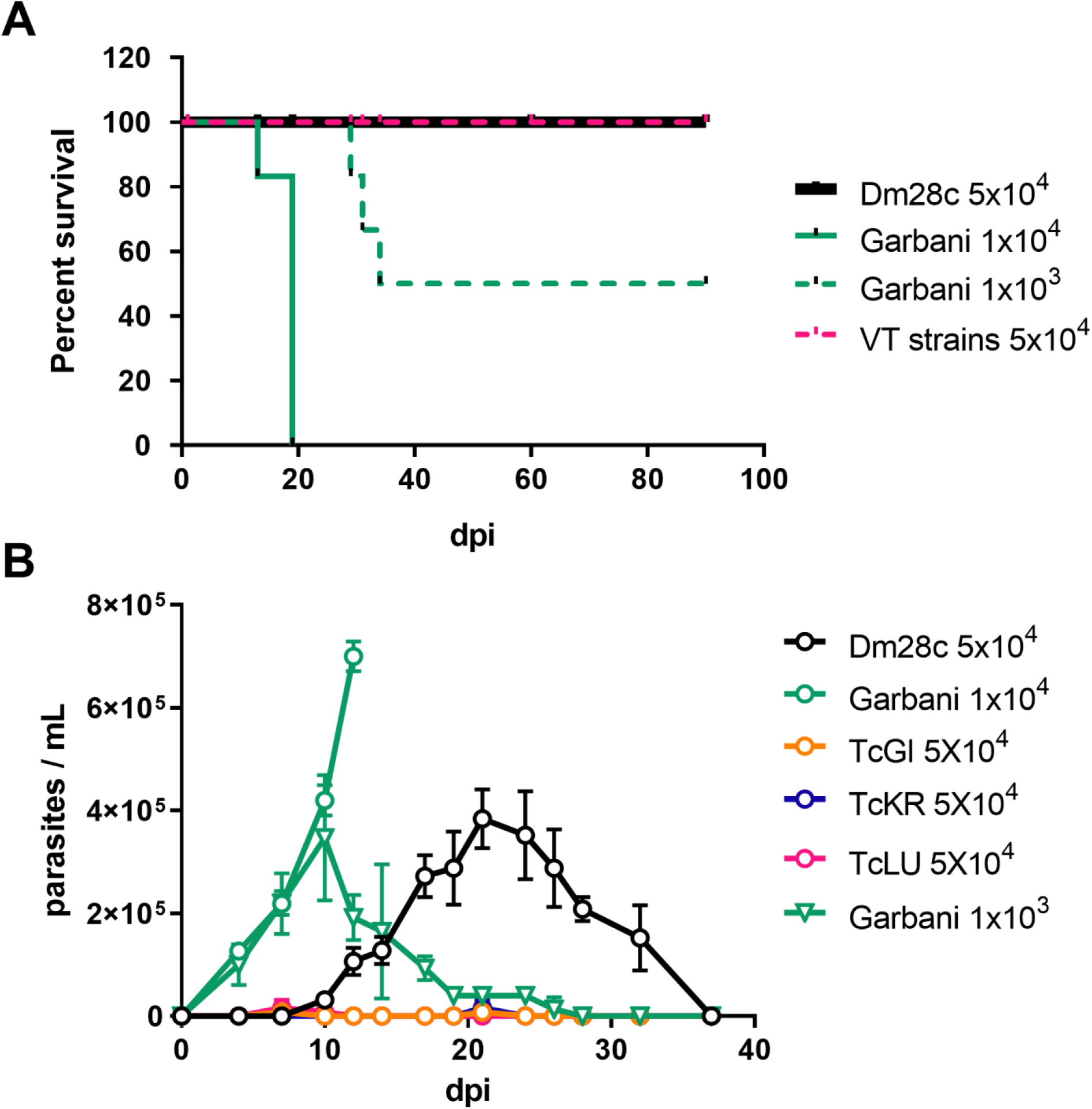
(A) Survival curves of mice infected with Dm28c (MV), Garbani (HV) and VT strains with the indicated initial inoculums. (B) Parasitemia along the course of infection of the different strains with the indicated inoculums.

### 3.3. Blood trypomastigotes of congenital Chagas isolates are unable to perform *in vitro* epimastigogenesis

Epimastigogenesis capacity was evaluated according to Kessler et al. (2017). After stress, a day-to-day follow-up was conducted to count epimastigotes and intermediate forms according to flagellar form, length and nucleus/kinetoplast position. As shown in **Figures 2A** and **2B**, at the start of the experiment, mostly trypomastigotes were observed and no intermediate forms were counted. By the 6^th^ day, approximately 60% of the population in the Dm28c and Garbani strains corresponded to epimastigote forms and 40% to intermediate forms, while the total population of parasites in VT isolates (TcGI, TcKR and TcLU) corresponded to intermediate forms. Finally, a stable 100% replicating epimastigote population was observed for the Dm28c and Garbani strains on the 10^th^ day of incubation but a different scenario of arrested intermediate forms was observed for the VT strains **(Figures 2A** and **2B** TcGi d2, d6 y d10). It is important to mention that differentiated Dm28c and Garbani epimastigotes were able to replicate and maintain a stable replicating population, whereas the cultures of VT strains (intermediate forms exclusively) were not able to replicate and died around the 20^th^ day. Attempts to maintain them in culture by changing media and adding supplements failed to induce replication or rescue the population.

**Figure 2.**
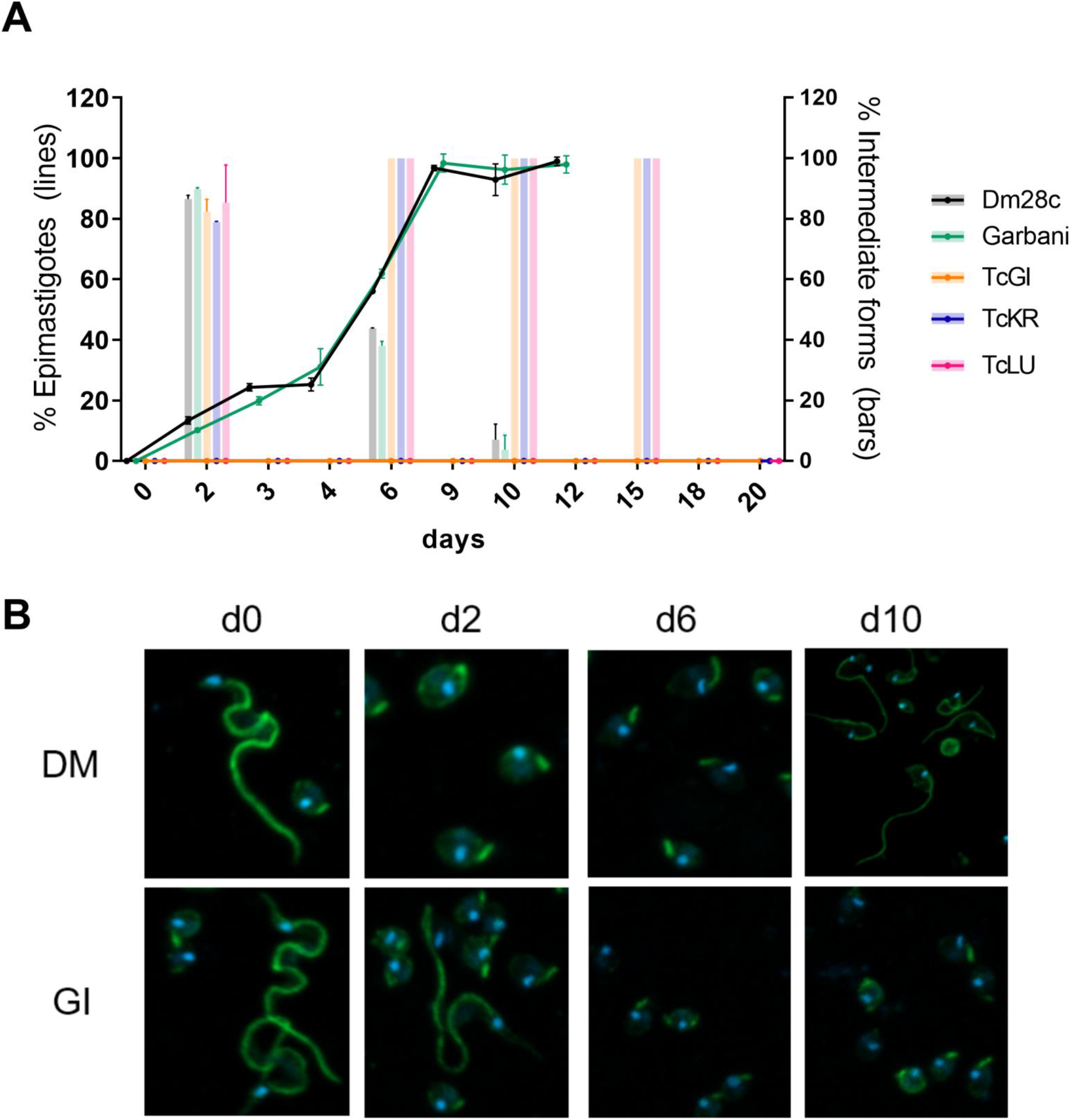
(A) Proportion of epimastigotes (lines) and intermediate forms (bars) out of total population of trypomastigotes during the epimastigogenesis experiment that lasted 20 days. (B) Immunofluorescence of the epimastigogenesis process of two strains Dm28c and TcGi (VT strain) that does not differentiate over time.

### 3.4. VT isolates are able to colonize different organs and induce a systemic immune response, similarly to the highly and moderately virulent strains

At day 30 pi, parasite DNA was detected in all organs regardless of the strain, and no statistically significant differences were found (**Figure 3A**). In gut tissues similar loads were observed for all the strains, but with higher values than those recorded for the rest of the tissues, suggesting all the strains in this particular model share a preferential tropism. As expected, when we observed mean values, parasite loads diminished almost to the non-infected control level (line in 1) in all tissues and for all strains after day 60 pi (**Figure 3B**). Dm28c infected organs such as the cardiac muscle, gut and uterus showed positive mean values but differences with the other strains were not significant. Notably, the parasite load of the TcKr strains in the uterus did not change within 30 or 60 days pi.

**Figure 3.**
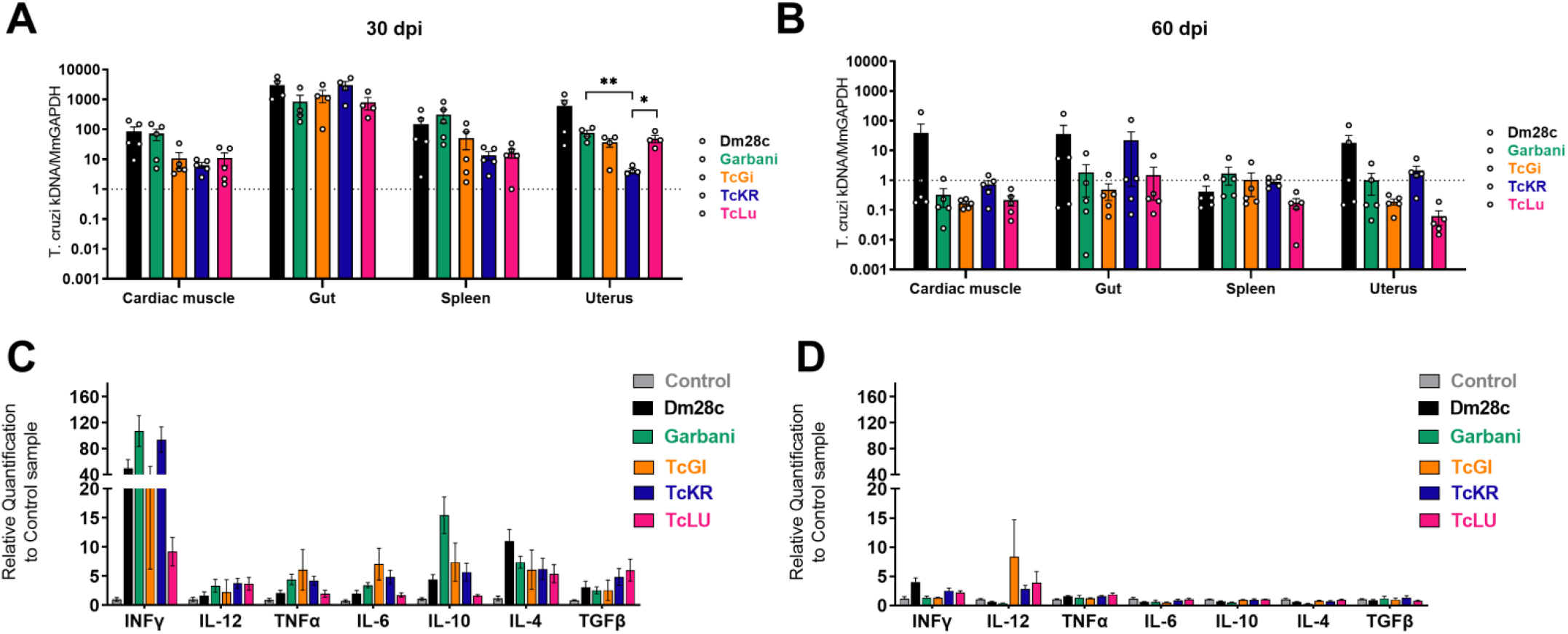
Parasitic loads in different tissues of mice infected with the different strains of *Trypanosoma cruzi* at 30 dpi (A) and 60 dpi (B). Level of expression of cytokines in splenic cells at 30 dpi (C) and 60 dpi (D).

The splenic response at day 30 pi measured by mRNA expression showed a strong upregulation of the different cytokines (IFNγ, IL12, TNF-α, IL6, IL10, IL4 and TGF-β) induced by infection with all parasite strains as shown in **Figure 3C**. All the cytokines evaluated were overexpressed in infected mice compared to uninfected control animals (grey bars). It is noteworthy that the general levels of each cytokine were comparable among all the strains regardless of the difference in virulence previously described. Garbani showed higher levels of expression for most genes than Dm28c, except for the anti-inflammatory cytokine IL4. A strong and more likely pro-inflammatory response was observed for the TcGi and TcLu strains whereas TcKr had lower levels of expression, except for TGFb. These results indicate that despite their low virulence, the VT strains reach the different organs and induce an immune response similar to that of the high and moderate virulence strains. At 60 days pi (**Figure 3D**), cytokine expression in infected groups fell to control levels, compatible with an ending of the acute phase, with the exception of IL12 levels that remained significantly upregulated in mice infected with the VT strains compared to the control group expression level, suggesting a long-lasting protective response.

### 3.5. Infections with the different strains do not significantly affect reproductive variables

To address whether mating, pregnancy and resorption rates were altered with infection and whether the strains were vertically transmitted to the offspring, a vertical transmission model was optimized **(Figure 4A**). As shown in **Figure 4B**, mating rates decreased with infection, while the mating rate of the control group was around 50%, infected groups had significantly lower rates: 40% for the group infected with the Garbani strain, and 20% for Dm28c, TcGI, TcKR and TcLU. Regarding pregnancy rates and resorption rates (measurement of implantation fate), no significant differences were observed for any of the groups compared to the control group. Altogether, these results indicate that under our conditions neither infection nor the strain has a negative impact on fertility or fetus viability.

**Figure 4.**
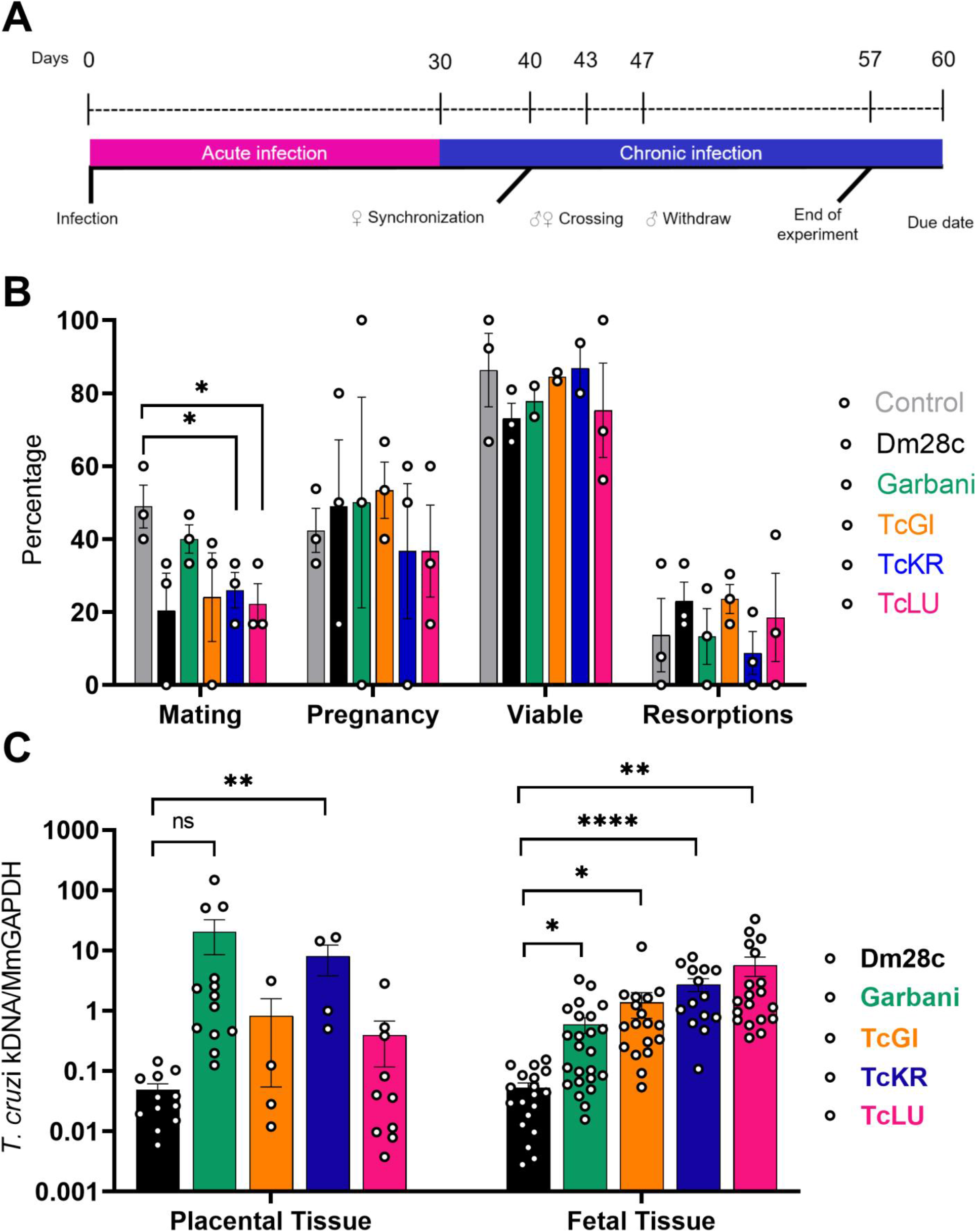
(A) Vertical transmission experiment scheme. (B) Mating, pregnancy, viable fetuses, and resorptions obtained from the vertical transmission experiment for the different experimental groups. (C) Parasitic loads determined in placenta and fetal tissue for the different experimental groups.

### 3.6. VT strains are more efficient than reference strains in transplacental transmission

After delivery, fetal, placental, uterine and resorption tissues were processed to detect parasite DNA. For each sample, an equivalent amount of tissue was processed from non-infected mice to set up a negativity threshold (1 in **Figure 4C**). A great heterogeneity was found among parasite loads in placental and fetal tissues as shown in **Figure 4C**. In the groups infected with the VT strains no parasite loads were found in the uterine tissue or the placenta (except for TcKR), and remarkably, all the groups infected with the VT strain showed a significantly higher parasite load in the fetal tissues, indicating that regardless of low virulence and parasite absence in the placenta, parasites were able to pass through the organ into the fetus more efficiently than the other (more virulent) strains, probably indicating transplacental passage early on gestation.

### 3.7. VT strains are more infective in human trophoblast derived cells compared to fibroblasts

After two hours of incubation, all the strains were able to infect human fibroblast as well as trophoblast cell lines. When using laboratory reference strains, the HFF line was more susceptible to infection by *T. cruzi* than the Swan cells, whereas the VT strains showed low infectivity in both cell lines; however, all of them infected Swan trophoblast cells more efficiently (**Figure 5A**). This can be better visualized by determining the Swan/HFF infection indexes: while the VT strain showed relative values of 4 and 10, values for the Garbani and Dm28c strains showed an inverted tendency with values below 1 (**Figure 5B**).

**Figure 5.**
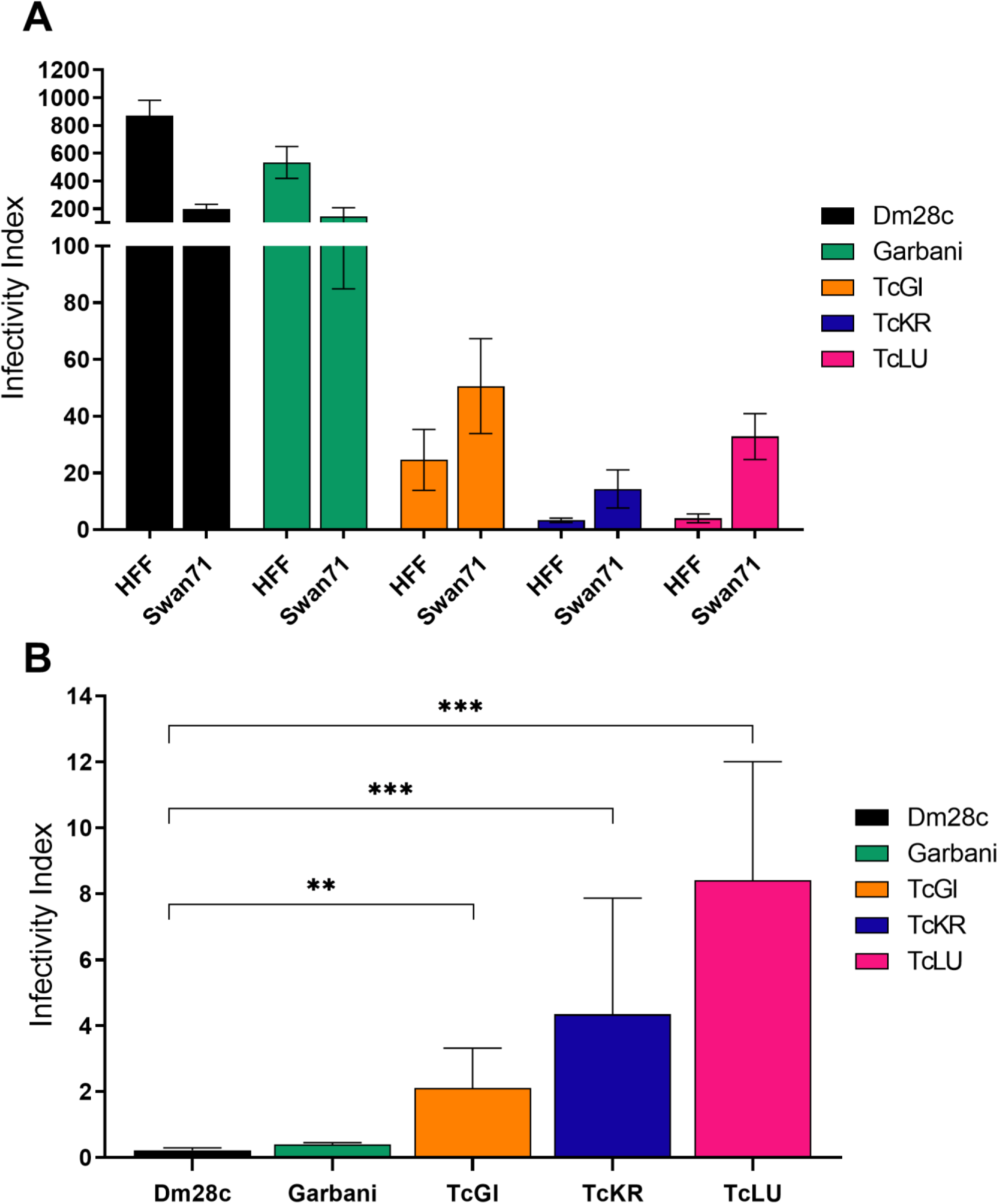
(A) Infectivity Index determined for each *T. cruzi* strain and in two different human cell lines, HFF (fibroblasts) and Swan71 (trophoblasts). (B) The relationship between Swan71 infection index respect to HFF infection index determined for the different strains.

### 3.8. VT isolates induce a unique pattern of placental gene expression

The effect of the different strains of *T. cruzi* on placentas was studied at the gene expression level by RNAseq. Taking into account the high variability of *in vivo* experiments, we carried out a PCA analysis whereby three groups were clearly identified (**Figure 6A**): i) Garbani, ii) Dm28c and non-infected mice, and iii) VT isolates, corresponding to high, medium and low virulence respectively. Their correlate in the hierarchical clustering of the differentially expressed genes (DEG) is shown in **Figure 6C**. With a cutoff of -1.5 ≥ Log2FC ≥ 1.5 and a *p*-value < 0.01, the highly virulent Garbani strain modulates the expression of 2507 genes (1114 up and 1393 down compared to the uninfected control condition), the VT strains modulate 408 genes (232 up and 176 down), and only two overexpressed genes were found in placentas of mice infected with Dm28c (**Figure 6C** and **Supplementary Table 3)**. The relation between up- and down-regulated genes is close to 1 for all conditions; none of the differentially expressed genes is shared by all the strains, and only 29 genes are shared by the Garbani and VT strains (8 up and 21 down) (**Figure 6B** and **supplementary table 4**). When the 30 most expressed genes (**Supplementary figure 2**) were compared with the most significant differentially expressed genes (**Supplementary figure 3**), their respective heatmaps were opposites of each other, in other words, placental responses to high- and low-virulence vertically transmitted strains were diametrically opposed. The same opposite images were obtained when gene ontology was analyzed (**Figure 7**). Notably, the most significant GO terms in the highly virulent strain were inflammation, cellular immune response and ribosomal proteins, all of them at the expense of over-expressed genes, while the most significant terms in the VT strains (mitosis, meiosis, cell cycle, DNA replication and response to DNA damage) consisted mainly of down-regulated genes. In addition, the processes enriched in one condition were not affected in the other. When we dissected each term into its components, a surprising number of up- or down-regulated genes were found as an exclusive response to each condition, that is, to the tropism of the strains. (**Supplementary Figure 4**). On the other hand, terms that were significantly down-regulated in Garbani were related to transport, lipid metabolism, cell morphology and adherence, whereas the up-regulated terms in the VT strains were import- and secretion-related genes which were down-regulated in the placentas of mice infected with the highly virulent strain.

**Figure 6.**
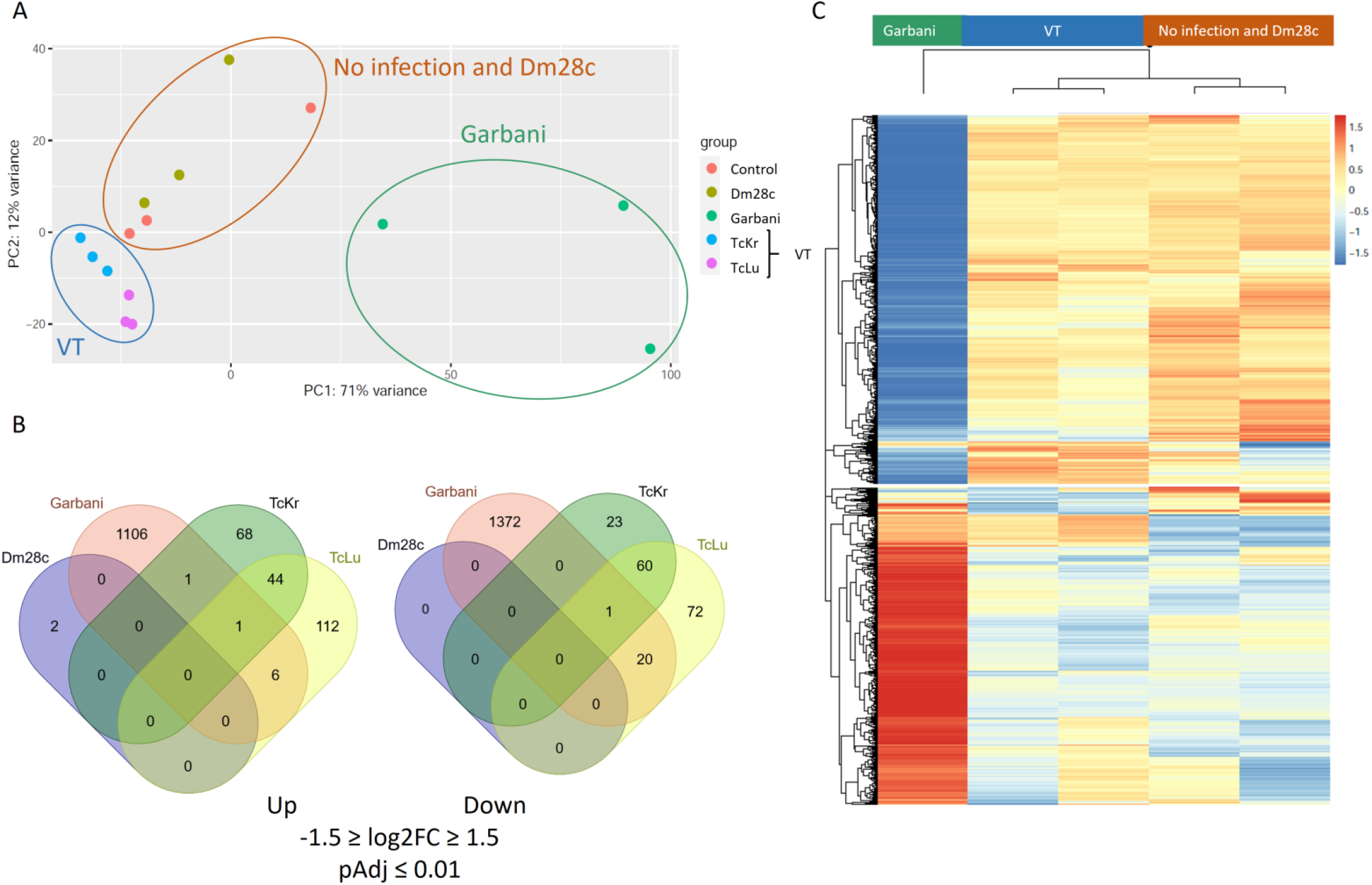
(A) Principal Component Analysis (PCA) of RNA sequencing of the different biological replicates of placentas. (B) Venn diagrams of differentially expressed genes (DEGs) -1.5 ≥ Log2FC ≥ 1.5 and Padj ≤ 0.01 in infected placentas compared to control placentas. (C) Hierarchical cluster of normalized read counts of DEGs. No infection (control placentas), Garbani (HV strain), Dm28c (MV strain), and TcKr and TcLu (VT strains).

**Figure 7.**
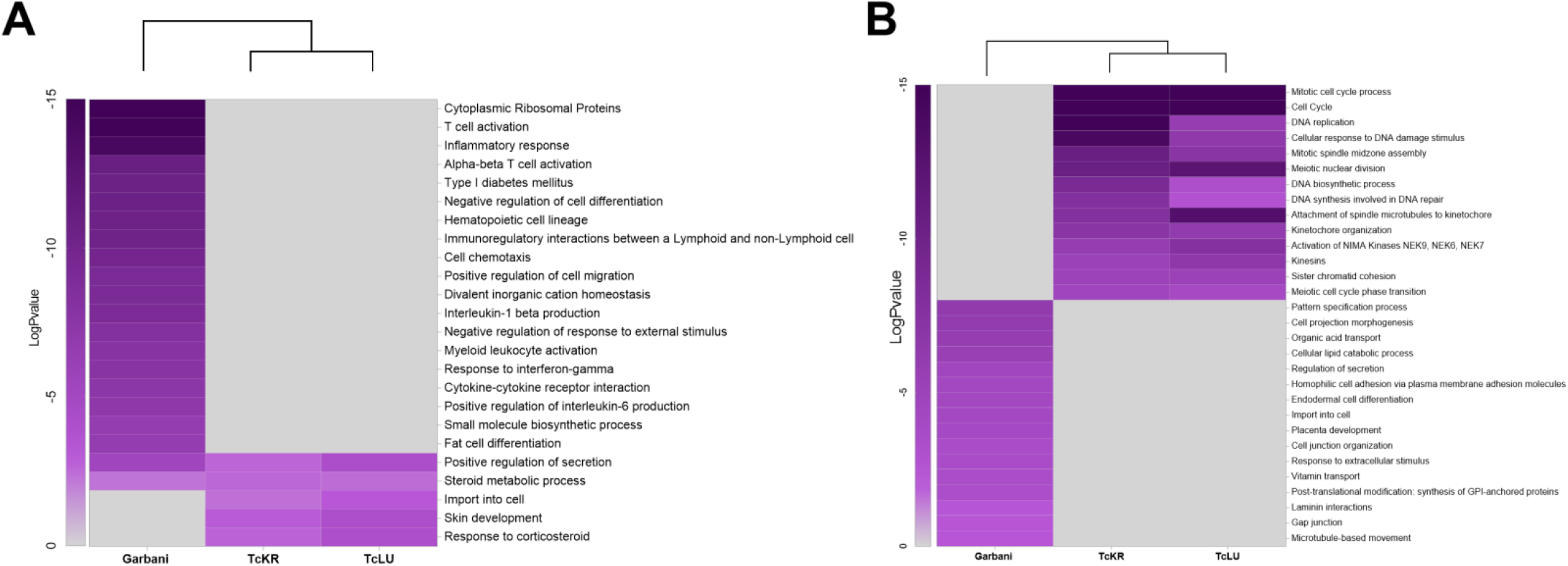
Heatmaps showing enriched GO terms among the up-regulated DEGs (A) and down-regulated DEGs (B) in placentas derived from mice infected with Garbani (HV), TcKr and TcLU (both VT strains).

Specific gene families exhibiting inverse regulation in the Garbani and VT placentas are shown in **Figure 8**. Among these groups, those worth mentioning are the ones playing important roles in the maternal-fetal interphase: immunomodulators such as genes encoding pregnancy-specific glycoproteins (*psg*), carcinoembryonic antigen cell adhesion molecules (*ceacam*), and prolactins (*prl*), in addition to those related to cell permeability and cell adhesion like solute carriers (*slc*), sodium channels (*scn*), desmogleins (*dsg*), and claudins (*cld*). Differential regulation among the Garbani and VT isolates also involved matrix metalloproteinases (*mmp*) and the adamlysin family (*adam*) and their intracellular (*timp*) and secreted (*sfrp*) inhibitors. Gene families involved in defense mechanisms showed opposite modulation in the different conditions, for instance, metallothioneins (*Mt*), proteins that sequester metals, were up-regulated in the Garbani group, signaling lymphocyte activation molecules (*slam*), and so were inflammosome-related factors.

**Figure 8.**
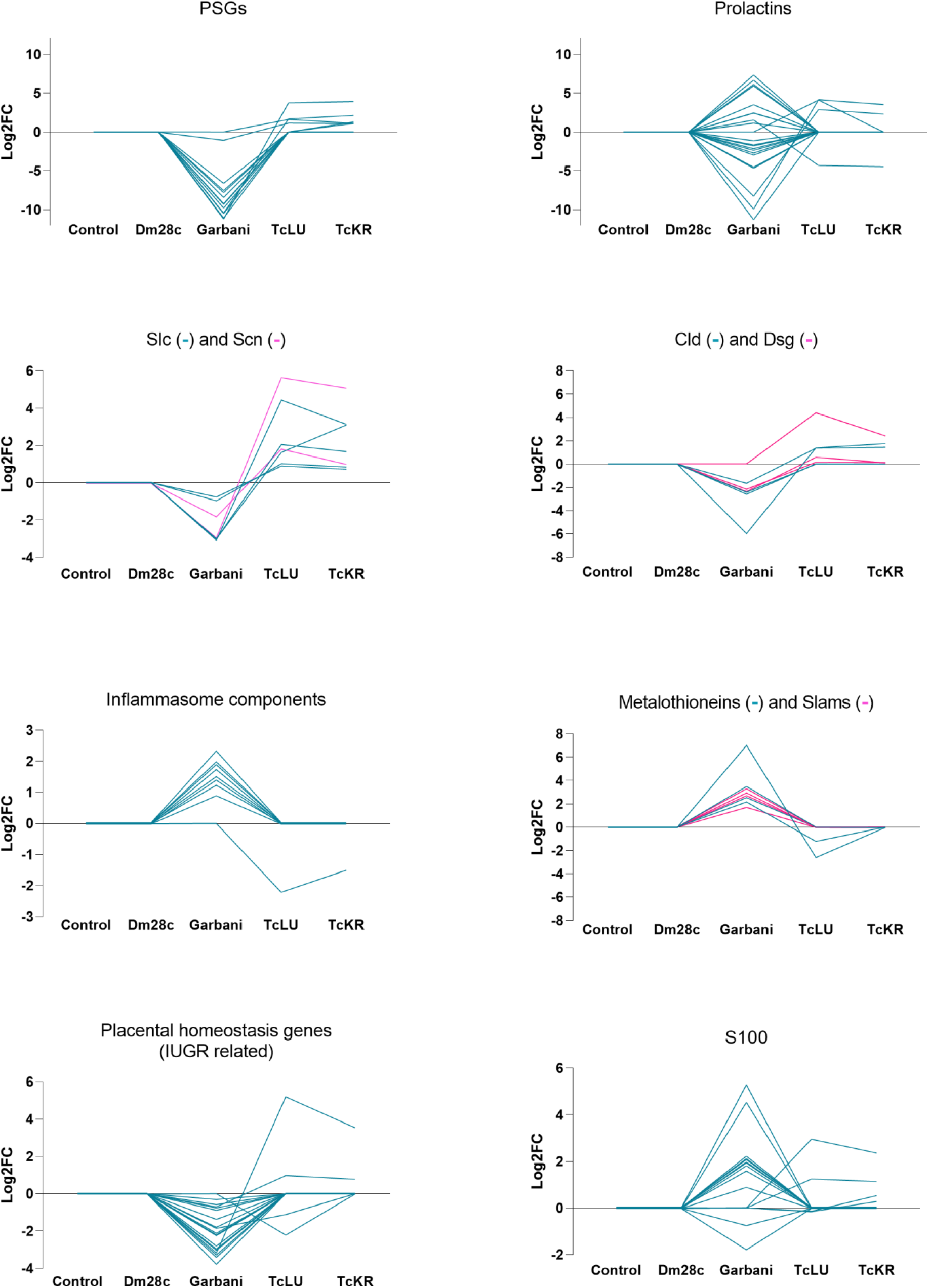
Differential expression (log2FC) found in groups of genes of interest comparative between the different experimental groups.

In summary, the three groups found concerning *T. cruzi* and placental relation/tropism also had differential placental responses evidenced in the RNAseq experiment: while placentas of the groups infected with the medium virulence strain resembled control placentas (and in fact, transmission did not occur), the high- and low-virulence groups exhibited completely different and opposite responses.

### 3.9. The molecular footprint of placentas correlates with histopathological alterations

Results from the analysis of placental sections stained with hematoxylin and eosin indicated that placentas of the Garbani group were the most affected, showing a high score in edema, inflammation, necrosis and tissue degeneration. As shown in **Figure 9**, pathological alterations were found in both the maternal decidua basalis and the labyrinth zone of placentas of mice infected with the Garbani strain. Regarding edema and inflammation scores, the Garbani strain tops the rank, followed by Dm28c, and last, TcLU with mild inflammation. TcKR placentas do not show inflammation, degeneration or necrosis. It is important to mention that no amastigote nests were identified in any of the samples using this staining method.

**Figure 9.**
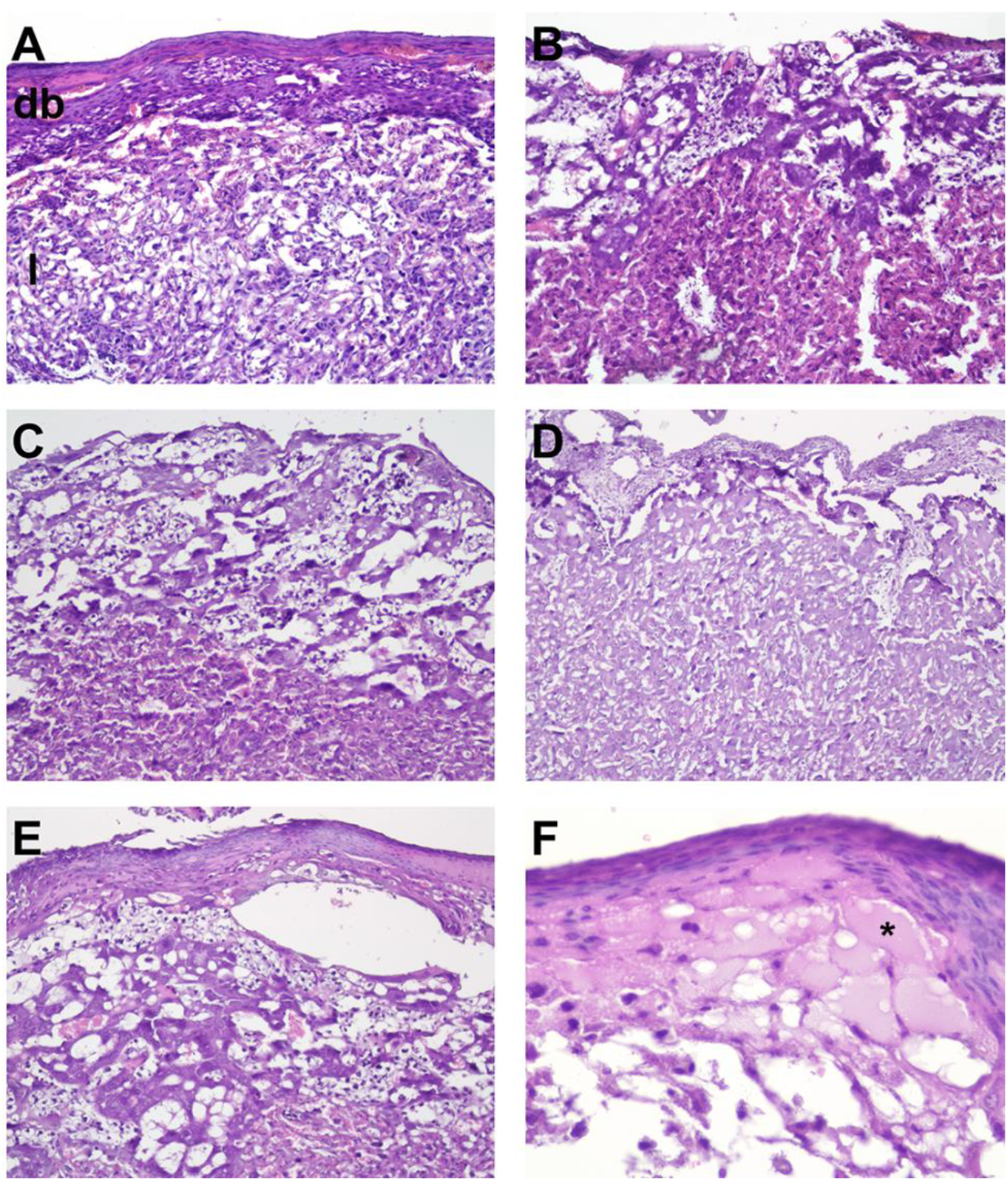
Histopathological analysis of placentas. Note that in all panel figures, the side of the placenta facing the mother (maternal side) is at the top and the side facing the fetus (fetal side) is at the bottom. Maternal decidua basalis (db) and fetal labyrinth (l) are shown in (A) (Control, H&E, 10x). (B) Mild edema of decidua basalis (DM28c, H&E, 10x). (C) Mild edema of decidua basalis (TcLu, H&E, 10x). (D) Moderate edema of decidua basalis and labyrinth (TcKR, H&E, 10x). (E) Moderate edema of decidua basalis and labyrinth (TcGa, H&E, 10x). (F) Detail of moderate edema of decidua basalis (asterisk) and necrosis with mild mononuclear inflammatory infiltrate of labyrinth (arrowhead) (TcGa, H&E, 40x).

## 4. Discussion

This work began with clinical observation. In the majority of congenital Chagas disease cases, the mothers are diagnosed during pregnancy at routine controls, due to the absence of symptoms and familial clustering (several cases of Congenital Chagas among one family cluster) is frequently observed (Altcheh et al., 2005; Muñoz et al., 2009; Howard et al., 2014). In this context, our working hypothesis was that there are *T. cruzi* strains adapted and specialized to vertical transmission that probably differ substantially from commonly used laboratory strains. We selected the particular scenario of a formerly endemic region where vectorial transmission has been uninterrupted for at least 25 years (Salvatella, 2016), and focused on babies born with the following conditions: i) Chagas Disease detected in the mother during routine pregnancy studies, ii) asymptomatic mother, iii) familial clustering, and iv) *T. cruzi* vertically transmitted to the baby, from which we obtained isolates. Throughout this work these isolates are generically named VT (for vertical transmission), whereas the reference strains of high (Garbani) and medium (Dm28c) virulence will from now on be referred to as HV and MV, respectively.

The first question was whether mice infected with VT isolates would mimic human-like clinical behavior and outcome i.e., low virulence, undetectable parasitemias, and classical *T. cruzi* persistence strategies. Both the survival curves and the persistence tests confirmed that they had very low virulence. Likewise, we determined in the *in vivo* assays that Garbani and Dm28c are strains of high and moderate virulence, which makes them suitable controls (**Figure 1**). VT strains, however, showed persistence capacity since they were able to infect all organs studied (heart, gut, spleen, uterus; **Figure 3**) and induce a systemic immune response comparable to that observed during infection with the HV and MV strains. *T. cruzi* infection in Balbc/J induced a Th1 response (Ferreira et al., 2018) and we found that in all cases INF*γ*, IL12, TNF*α* and IL6 were up-regulated, as well as the anti-inflammatory cytokines like IL10 and IL4, without significant differences between strains. Both parasite clearance from organs and control levels of cytokine expression were reached by day 60 pi in mice belonging to the different strain infected groups (**Figure 3**). These results clearly show that VT parasites harbor intrinsic characteristics that allow them to disseminate and induce a competent immune response but are virtually non-detectable in blood. We can assert that the murine model reproduces the characteristics of low virulence observed in humans and that we have two adequate controls (medium and high virulence strains) for subsequent comparative studies.

The second question was to evaluate the impact of VT strain infection on reproductive and vertical transmission parameters **(Figure 4)**. We developed a murine model of mating and pregnancy in mice and found that *T. cruzi* infection affected mating in all cases, regardless of the responsible strain; although not due to virulence, since the HV strain showed the highest mating percentage. On the contrary, the infection did not affect pregnancy or spontaneous abortion rates since differences observed in these parameters were not significant. It is worth mentioning that time of infection is an important factor for pregnancy outcome in female mice, since going through the acute phase close after embryo implantation induces unhealthy pregnancy outcomes (Cencig et al., 2013; Leon et al., 2017). Despite the effect on mating, we did not observe any impact on pregnancy rates in our model and our results are in accord with other reports (Cabeza Meckert et al., 1980; Solana et al., 2002). The finding that parasitic loads are higher in fetal tissue of VT infected groups than in fetal tissue of reference infected groups leads us to indicate that VT strains are specialized in vertical transmission (Figure 4C). Moreover, VT strains exhibited a more efficient persistence strategy since they colonized different organs and induced an immune response without generating major damage in the host (this was demonstrated in our murine model and clinical data indicates that this is also valid in humans). The VT strain transmission mode constitutes an example of highly successful parasitism which additionally dispenses of the insect vector. Concerning the reference strains, there were two completely different situations: **a)** the MV infected group showed the highest parasitic load in the uterus and the lowest signal in the placenta, and was not transmitted vertically at all; **b)** the HV infected group was vertically transmitted but with lower efficiency (low signal in fetal tissue and less infected individuals in the offspring) than the VT strains and showed low parasitic load in the uterus and a positive signal in the placenta. The inability of the MV strain to transmit vertically is a phenomenon that requires attention. Although it is beyond the scope of this work, we want to point out that this finding opens a question for future investigation: what is present or missing in the Dm28c strain that prevents it from being vertically transmitted? On the other hand, the HV strain capacity for vertical transmission leads us to postulate that it is due to high virulence at the cost of infecting almost any tissue and organ, which does not imply a specialized mechanism or placental tropism. In addition to being an example of inefficient parasitism, this type of strain, widely used in laboratories, does not seem to resemble the behavior expected for circulating strains in humans. Moreover, the observed placental tropism of VT strains has a correlation in an *in vitro* human model: a comparison of the Swan (trophoblast derived) index versus the HFF index can be used to show that this relationship is inverted in VT strains. The Swan/HFF infection index is higher in the VT strains than in the MV and HV strains (**Figure 5**). Finally, to further explore *T. cruzi* adaptation capacity to the mammal side of the cell cycle (and why not the human side) we assayed *in vitro* epimastigogenesis. As suspected, whereas reference strains shifted toward epimastigotes in the expected period, VT strains remained in intermediate form and never replicated (**Figure 2**). This process, which occurs naturally in the digestive tract of triatomine vectors, can be reproduced both *in vivo* (Navarro A. Angi S.; et al., 2012) and *in vitro* (Rondinelli et al., 1988; Kessler et al., 2017). Given that *in vivo* epimastigogenesis is only partially modelled by the *in vitro* process and that VT strains were able to replicate in triatomines from which they were isolated, our results suggest that parasite plasticity allows specialization and adaptation to non-vectorial transmission. Similar observations were made in other trypanosomatids, such as African trypanosomes that were mechanically transmitted between cattle during their introduction to America (in the absence of their natural vector, the tsetse fly), and selection occurred towards parasites specialized in the mammalian cycle, including the loss of guide RNAs necessary for complete mitochondrial gene editing (Greif et al., 2015). Future analysis of comparative genomics and gene profiling on VT strains is needed to dissect parasite molecular mechanisms involved in the adaptation to human infection through vertical transmission.

Our next question was: Are there changes at the placental gene expression level that could explain the specialization of VT strains to vertical transmission? We performed RNAseq on placentas of mice groups infected with the different strains and the main results can be observed on the PCA graph with the presence of three groups (**Figure 6A**). The hierarchical clustering of differentially expressed genes (**Figure 6C**) confirmed that HV and VT strains induce a totally different placental response. First, out of approximately 3,000 differentially expressed genes, most of them (∼87%) corresponded to placentas of the HV infected group, ∼13% to VT strains, and only 2 genes were found up-regulated in the Dm28c (MV) strain. Remarkably, none of the differentially expressed genes was shared by all strains, and only 29 genes were shared by the HV and VT strains. On one hand, the MV strain is unable to vertically transmit at all. Among those strains that are transmitted, there are two scenarios: while the HV strain induces dramatic changes in placental gene expression, mainly involving up-regulation of genes related to inflammation, VT strains do not affect immune response-related genes, but down-regulate genes belonging to mitosis, meiosis and cell cycle process probably inducing an arrested state in trophoblasts and other cell types present in the placenta. HV’s most significant changes involve gene induction and GO terms related to immune response, inflammatory processes, and ribosomal proteins (see below). A gene-by-gene heat map overview showed that placental responses resemble opposite images, where genes up-regulated in Garbani (HV) placentas are down-regulated or unchanged in VT placentas and vice versa.

As mentioned, one of the main differences among placental responses of the high virulent strain and VT infected placentas belong to immune and inflammatory processes (**Supplementary figure 4**). In addition, analysis of up-regulated genes belonging to the immune response GO term in VT strains showed that all were related to tolerance and anti-inflammatory processes, being unchanged or down-regulated in HV infected placentas (**blue frame in Supplementary figure 4**). Among them, it is worth mentioning *s100a14*, a gene that was found to be down-regulated in the placentas of Chagas seropositive mothers (Juiz et al., 2018). The protein encoded by this gene is known to induce the *mmp2* gene (matrix metalloproteinase 2) (Chen et al., 2012; Qian et al., 2016), which we found did not change in any conditions and is capable of inducing macrophage migration (Chen et al., 2015). Its up-regulation has also been linked to a condition of placental tissue inflammation called histologic chorioamnionitis (Kim et al., 2014). Some of the S100 proteins function via the ‘nutritional immunity’ mechanism by sequestration of essential metals required for intracellular parasites to thrive (Kozlyuk et al., 2019). Remarkably, eight other *s100* genes that react to Damage Associated Molecular Patterns (DAMPs) (*s100a1, s100a3, s100a4, s100a6, s100a7, s100a9, s100a10* and *s100a11*) were up-regulated in HV and not in VT infected placentas. The fact that these eight genes are RAGE (Receptor for Advanced Glycation Endproducts) dependent is a fundamental difference with *S100A14* (down-regulated in HV), whose activation is RAGE independent. Since RAGE is a pro-inflammatory pathway, the activation of RAGE independent defense mechanisms may be attenuated in an anti-inflammatory environment like those found in VT placentas.

Metallothionein genes *mt1, mt2, mt3* and *mt4* were up-regulated in the HV group and down-regulated or unchanged in VT groups **(Figure 8)**. Their gene products were cytosolic proteins that mitigate metal poisoning and alleviate superoxide stress (Subramanian Vignesh and Deepe Jr., 2017). Inflammasome-associated cytokines have recently been described to be constitutively released from healthy pregnancy placentas. This state is considered an immunological signature of health, which is far from the past belief of an immunosuppressive state (Megli and Coyne, 2021). We found major inflammasome components differentially regulated in the HV-infected placentas, which suggests an unhealthy state that does not occur in VT infected placentas. Inflammasome components *nlrp1, nlrp3, pycard, gsdmd, casp1, casp4* and *casp12* were up-regulated in the placentas of HV and unchanged in the VT infected group (**Figure 8**). Up-regulation of inflammasome genes has been described (Megli et al., 2021) during placental infection with *Listeria monocytogenes* (a bacteria associated with poor pregnancy outcomes), similar to what we found for the HV strain but not in VT placentas.

The pregnancy-related genes (PSGs) with important roles in the maternal-fetal interphase were found to be differentially modulated between the two trans-placental passage models. PSGs are the most abundant trophoblastic proteins in maternal blood during pregnancy. They have immunomodulatory, pro-angiogenic and anti-platelet functions and regulate maternal fetal interactions (Moore and Dveksler, 2014). Low levels of PSGs (observed in HV, but not in VT placentas) are associated with fetal growth restrictions (Moore and Dveksler, 2014). The fact that 16 PSGs were down-regulated in placentas infected with HV and unchanged in VT placentas (**Figure 8**), correlates with the healthier state of VT placentas, the impairment of HV placentas to control immune responses, and the fact that low birth weight is observed for some babies born with congenital Chagas (Torrico et al., 2004). Moreover, some of these pregnancy-related genes have been directly implicated in the regulation of microbial infections. Prolactin for instance, mainly modulated in placentas of the HV group, has been proposed as an immunomodulatory molecule that enhances *Toxocara canis* migration to the uterus and mammary glands, allowing vertical transmission (Rio-Araiza et al., 2018). High prolactin serum levels have been associated with seronegativity to *Toxoplasma gondii* (Mohammadpour et al., 2019), which is capable of directly binding to the parasite and restricting intracellular growth and competent infection (reviewed in (Galvan-Ramirez Mde et al., 2014)). Another pregnancy-related protein is pregnancy zone protein (PZP), an alpha macroglobulin that is highly expressed during late pregnancy and has the capacity to inhibit proteinases. PZP levels increase during the course of acute Chagas in children and reportedly interact with *T. cruzi*’s cruzipain, the parasite’s main proteinase, inhibiting proteolytic capacity in order to minimize harm (Ramos et al., 2002). In our experiments, PZP was down-regulated in HV placentas and unchanged in VT. Regulation patterns of these pregnancy-related genes obtained in this work support the differential modulation of our models, evidencing that the passage of VT strains does not interfere with healthy placental function, therefore allowing silent transplacental passage.

It is worth mentioning a group of genes, related to membrane permeability and trafficking as they are sodium channels and solute carriers, which were up-regulated in placentas of the VT infected group, while down-regulated or unchanged in the HV group: *Scn9a, Slc22a13, Slc9b2, Slc6a1, Slc17a2* (**Figure 8**). Some of these genes have been previously associated with HIV infection and immunodeficiency, and were modulated by HCV in favor of viral propagation and establishment of chronic infection (Nguyen et al., 2018). Specifically for *T. cruzi* placental infection, several other solute carriers were found to be up-regulated in experimental infections (Castillo et al., 2018). Parasites are known to regulate cell-cell adhesions as part of their pathogenic mechanisms specifically by inhibiting gene expression and debilitating unions (reviewed in (Guttman and Finlay, 2009)). A set of genes, key components of cell-cell and cell-matrix unions that confer tissue stability and prevent paracellular passage of pathogens (Jimenez-Pelayo et al., 2017), were up-regulated in the VT infected placentas and down-regulated or unchanged in HV placentas. In particular, *Cldn* and *Dsg genes* are noteworthy, since they are components of tight junctions and desmosomes respectively, both related to paracellular metabolite passage and conductance (**Figure 8**). Additionally, *Ocln*, another component of tight junctions, was up-regulated with TcKR transplacental passage. Taken together, these results indicate that in addition to an inflammatory response, the HV strain induces changes that favor paracellular passage, already described for several pathogens (Guttman and Finlay 2009), including *T. cruzi* (Rodriguez et al., 2020). In opposition, VT strains induce junction-related genes, which should favor intracellular placental passage, contributing to immune escape. This type of strategy has been described for *Plasmodium* where, through a lysis-free mechanism of cell-to-cell passage called cell traversal, the parasite migrates through the tissue until it finds the proper microenvironment to settle and replicate (Yang et al., 2017). Transcriptomic results indicating tissue damage in placentas infected with the HV strains and more importantly, maintenance of tissue integrity in placentas of the VT infected groups, are supported by the histopathological analysis summarized in **Figure 9**. Finally, results obtained for VT strains, involving low replication and high placental tropism coupled with immunological silence without tissue damage, suggest that this strategic mechanism is exploited by specialized *T. cruzi* isolates and should be further studied. Generally, transcriptomic analysis found in the literature describes placental responses that resemble our results regarding the HV strain, concluding that parasite passage is more likely to be due to tissue damage and inflammation, which may lower parasite load but also allow passage.

In summary, we identified two different vertical transmission strategies that are represented in **Figure 10**:

**Figure 10.**
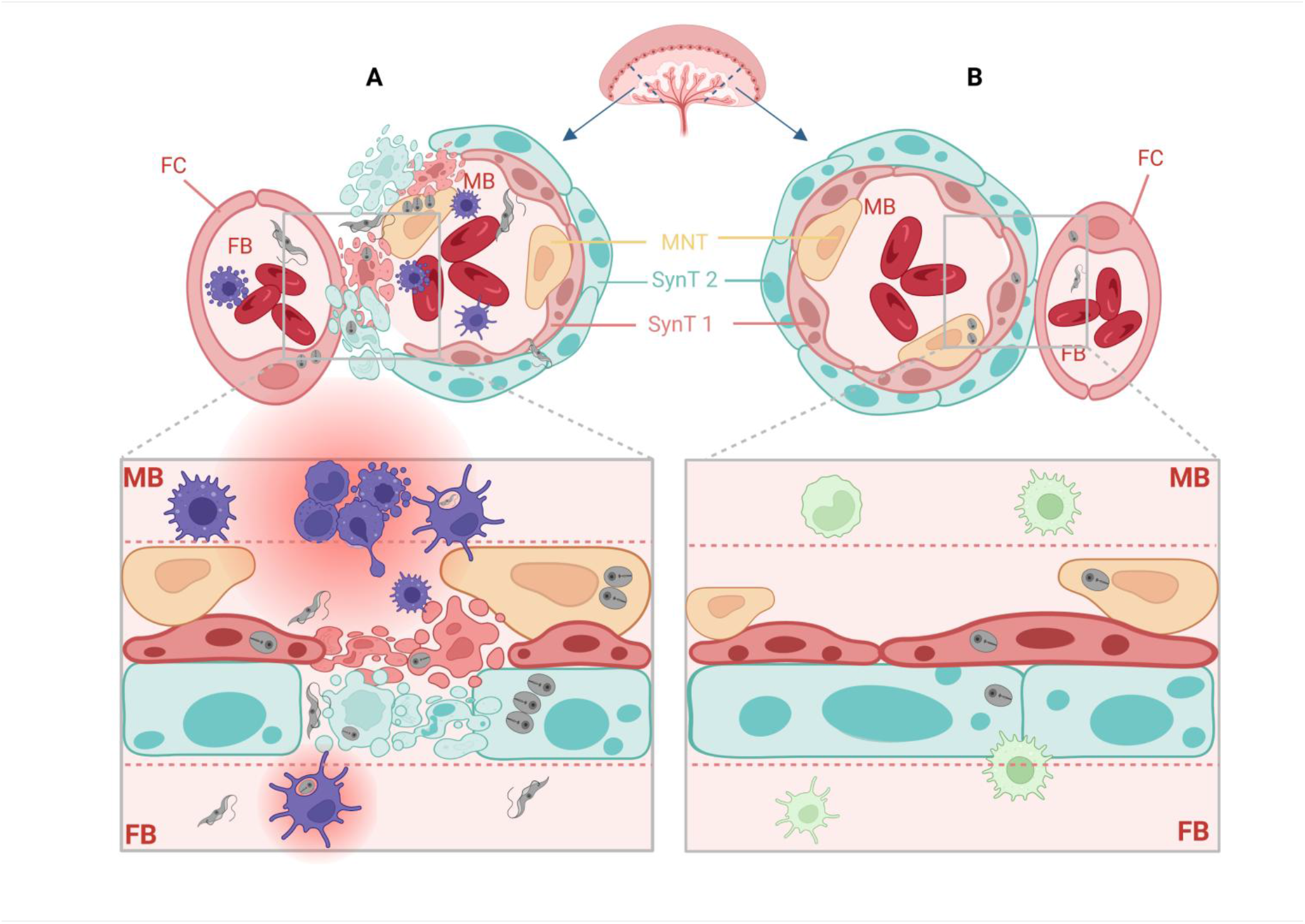
Two vertical transmission strategies at the murine maternal-fetal interphase. (A) Unspecific and without preferential tropism towards the placenta observed in the high virulent strain Garbani, *T. cruzi* is vertically transmitted inducing tissue damage and a strong pro-inflammatory placental response. (B) Specific and placentotropic strategy observed for VT isolates, *T. cruzi* is vertically transmitted in a silent manner, without causing damage and without inducing a pro-inflammatory response. Created with biorender.com.

1. Unspecific or non-preferential tropism toward the placenta. These strains (HV) are vertically transmitted inducing damage due to high virulence, evidenced by the pro-inflammatory placental response to infection. We think that this kind of virulent pattern does not necessarily correspond to what is seen in clinical cases of congenital Chagas, except for acute infection acquired during pregnancy which presents higher vertical transmission frequencies (Hermann et al., 2004; Moretti et al., 2005).
2. Specific and placentotropic. This is the case of VT isolates, which present mild immune response modulation and a characteristic profile or regulation involving small changes. The mechanism used by these strains constitutes a specialization to human infection and non-vectorial transmission of *T. cruzi*. It evidences an adaptative strategy to achieve vertical transmission without causing damage or illness that may jeopardize individual health. We postulate that this is the most common strategy in congenital Chagas, supported by the observation that in most of these cases the disease is diagnosed during pregnancy, since no signs or symptoms present. In addition, previous results regarding exacerbated immune responses, like the one we observed for Garbani (HV), are typical of laboratory virulent strains and biased because *in vitro* models fail to reproduce what happens in the maternal-fetal interphase.

## 5. Acknowledgments and funding

We thank Dr. Estela Bevilacqua for critical reading of this manuscript, Dr. Alejandro Schijman for confirming our DTU results, Dr. Gil Mor for kindly sharing the SWAN-71 cell line, Dr. Miriam Postan for kindly providing the Garbani strain, Dr. Guzmán Alvarez for Bzn and Nfx drugs, Dr. Sergio Schenkman for Mab25 antibody, and Grazzia Rey for helpful advice.

This work was supported by ERANET-LAC project ERANet17/HLH0142 (to Carlos Robello). PFT was a PEDECIBA student and received grants from Agencia Nacional de Investigación e Innovación (ANII, Uruguay) and Comisión Académica de Posgrado (Universidad de la República, Uruguay). PFT, GG, GL, AC, JMV and CR are members of the Sistema Nacional de Investigadores (SNI, ANII). GG, MC and CR are PEDECIBA researchers.

