## Supplementary figures and images for "*Trypanosoma cruzi* isolates naturally adapted to congenital transmission display a unique strategy of transplacental passage"

### Figure S1

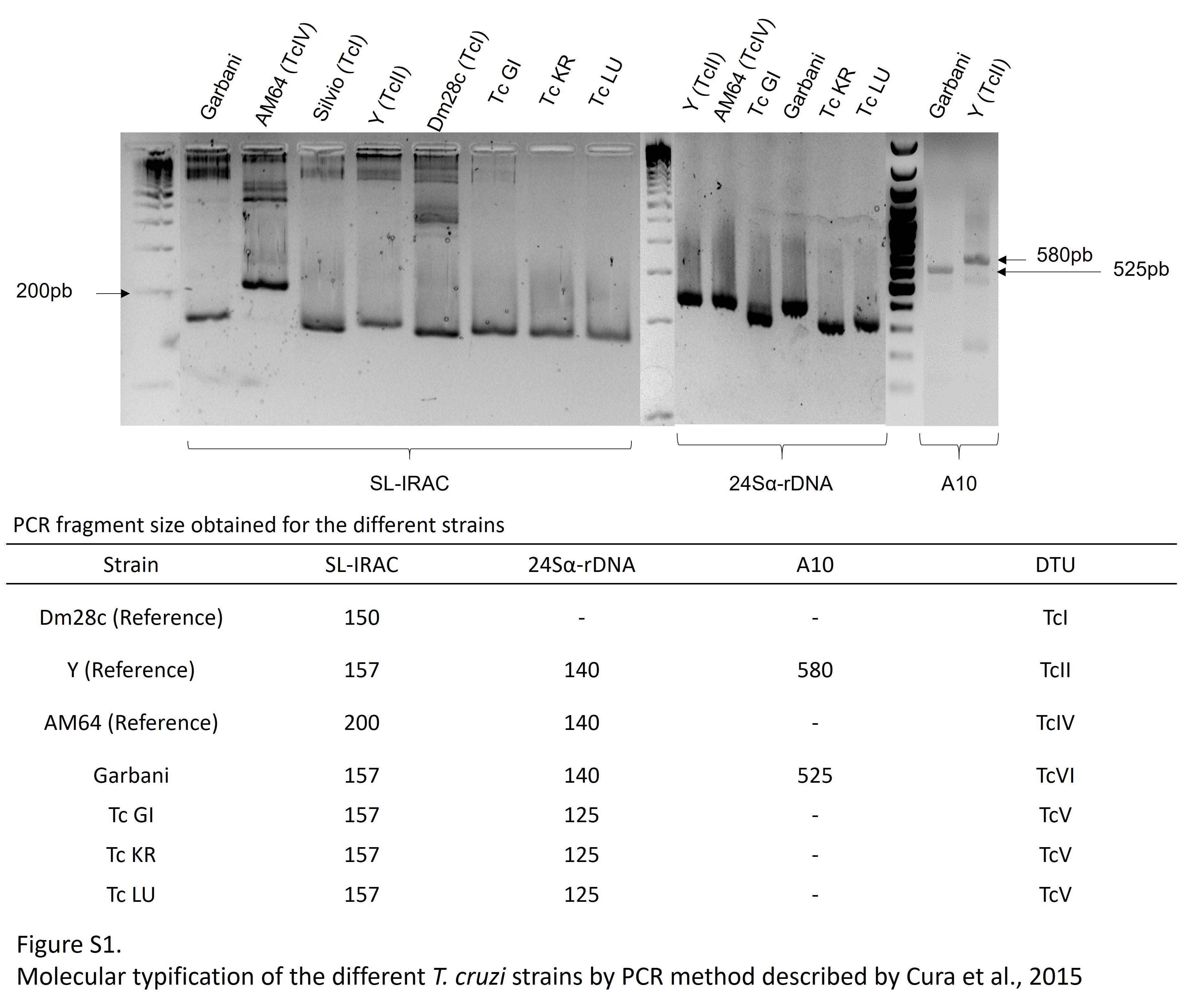

### File S2

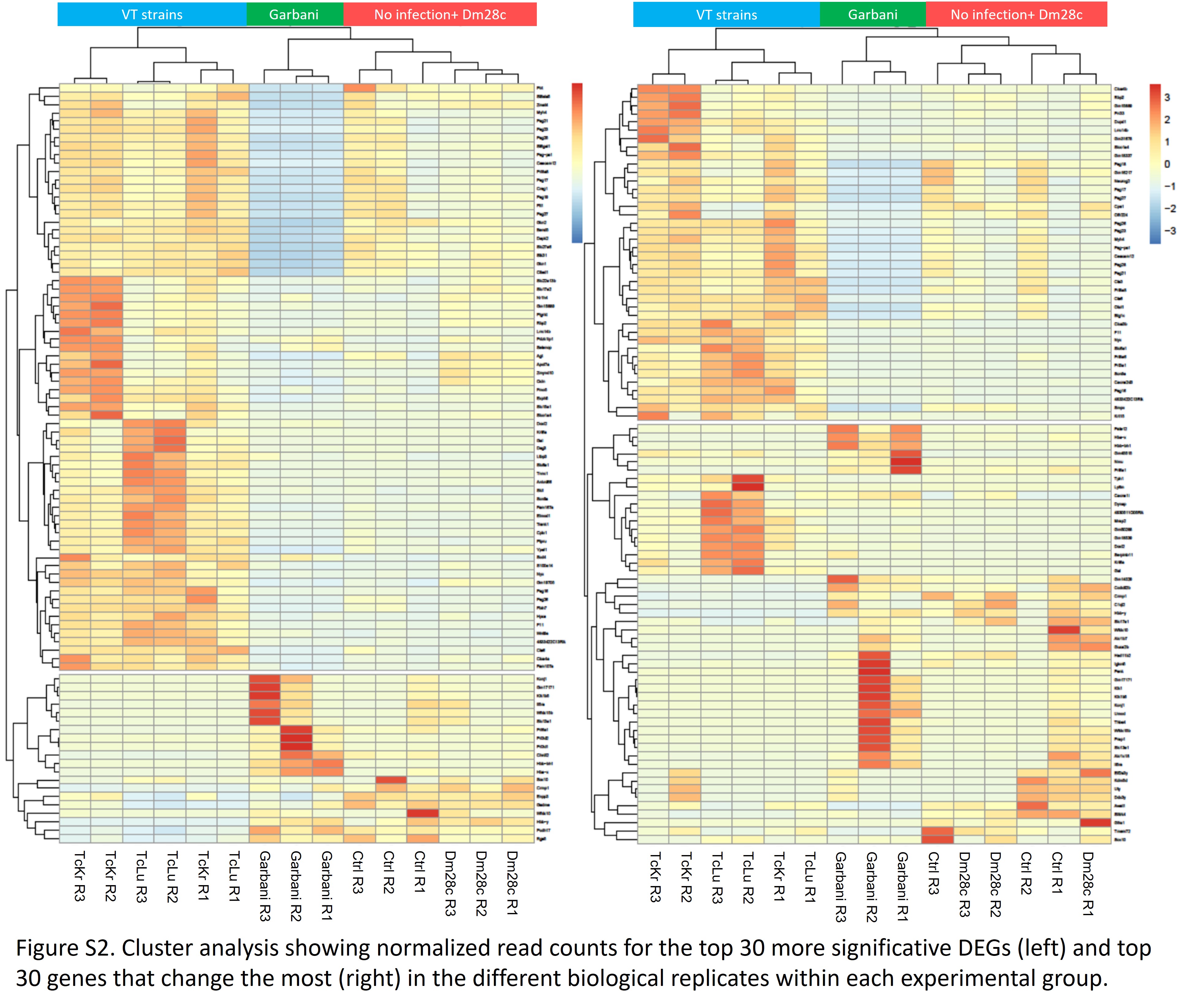

### File S3

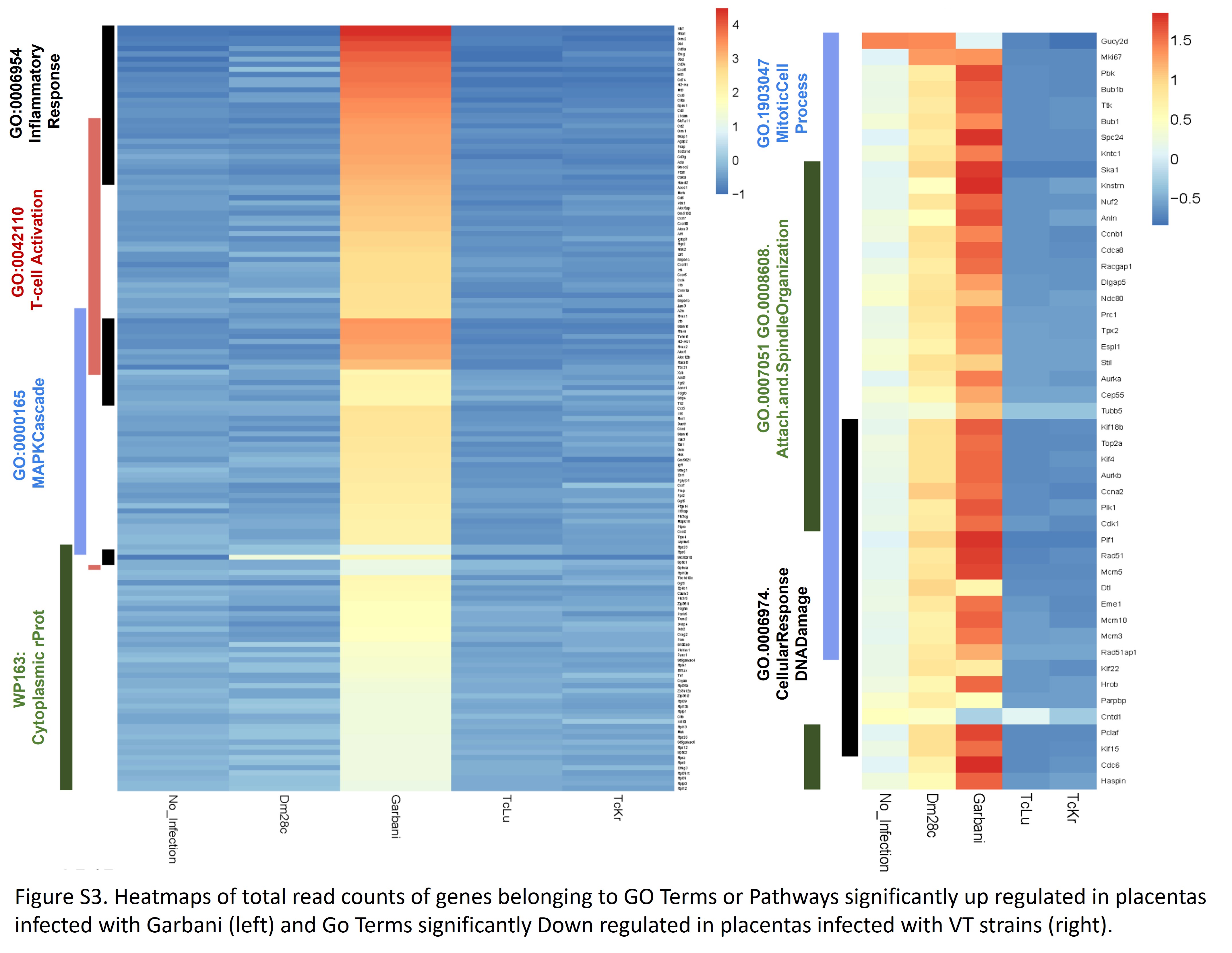
