## Supplementary material for "*Trypanosoma cruzi* isolates naturally adapted to congenital transmission display a unique strategy of transplacental passage": Figure S4

**Supplementary figure 4.**

**Heatmaps represent row scaled normalized mean (biological replicates) read count values for each strain infected placenta. Each heatmap shows the behavior of all genes from a selected pathway.**

1. **Immune response (blue box indicates genes related to tolerance and anti-inflammatory processes, upregulated in VT strains)**
2. **T-cell activation**
3. **Inflammatory response**
4. **Cell junctions**
5. **Cell cycle and other related**
6. **Mitotic cell cycle process**
7. **Cell projections and morphogenesis**
8. **Import to the cell**
9. **Positive regulation of secretion and import**
10. **Positive regulation of secretion**
11. **Ribosomal citoplasmatic proteins**

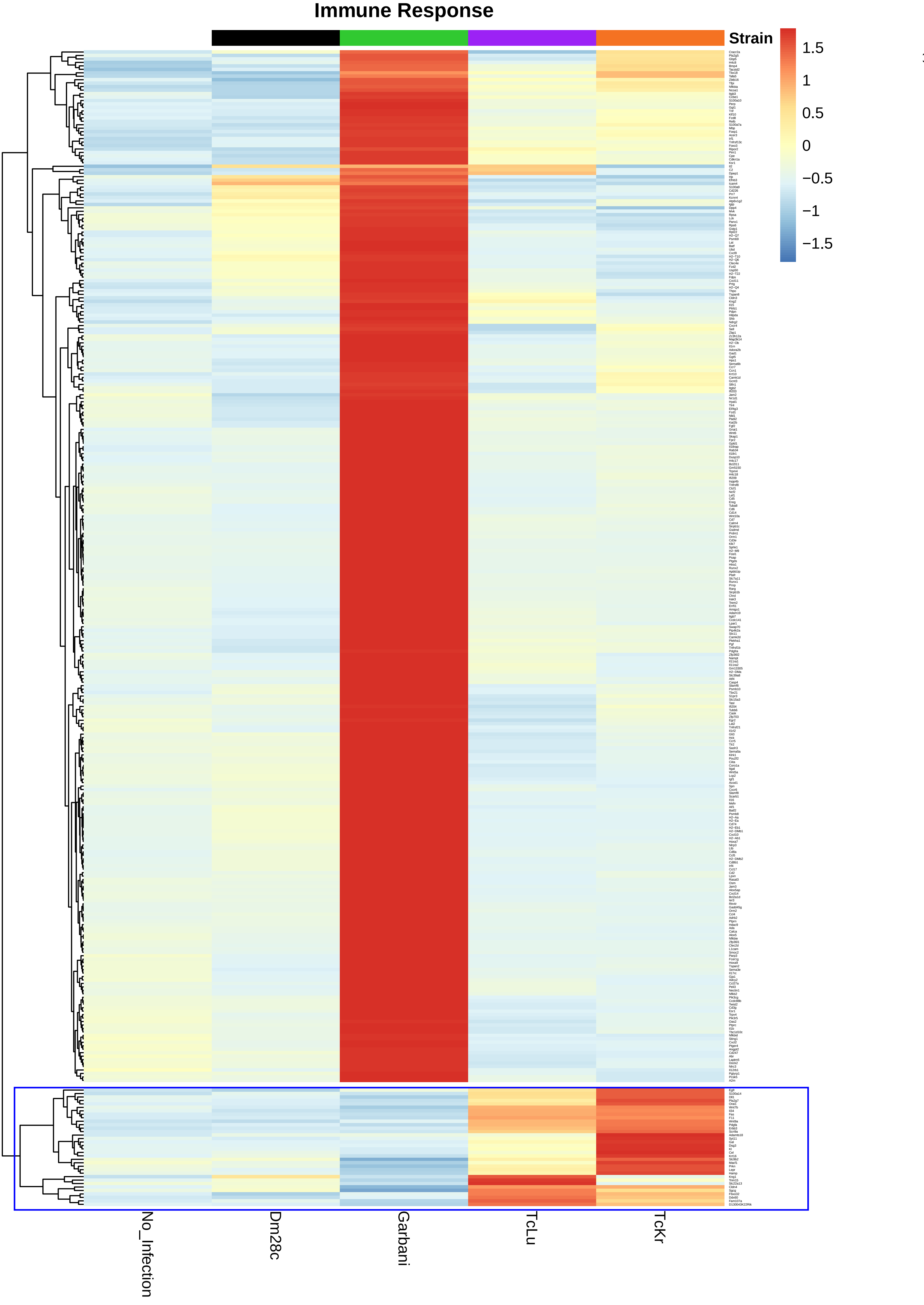
Immune Response
Strain
1.5
1
0.5
0
−0.5
−1
−1.5
No_Infection
Dm28c
Garbani
TcLu
TcKr

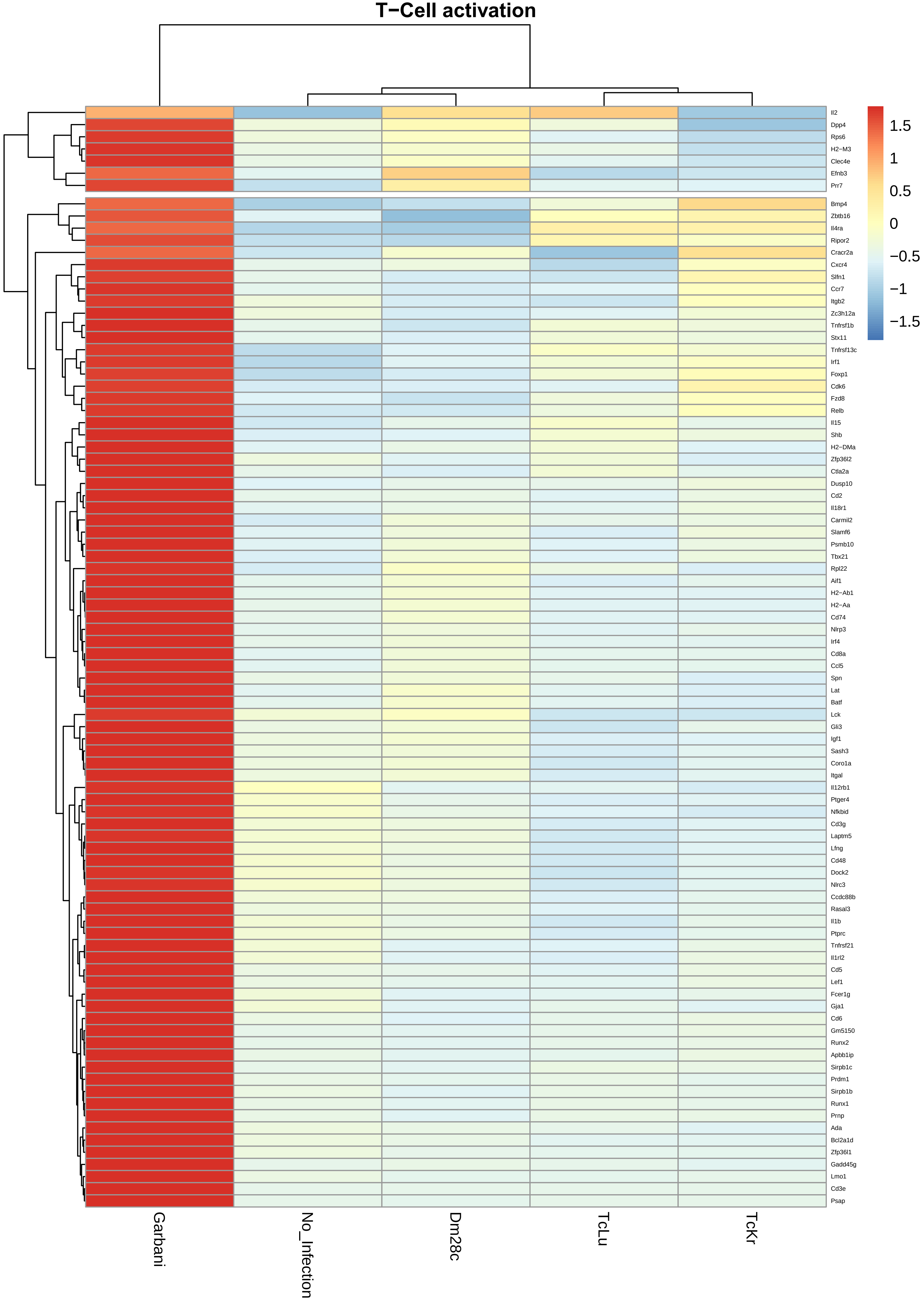
T−Cell activation
Il2
Dpp4
Rps6
H2−M3
Clec4e
Efnb3
Prr7
Bmp4
Zbtb16
Il4ra
Ripor2
Cracr2a
Cxcr4
Slfn1
Ccr7
Itgb2
Zc3h12a
Tnfrsf1b
Stx11
Tnfrsf13c
Irf1
Foxp1
Cdk6
Fzd8
Relb
Il15
Shb
H2−DMa
Zfp36l2
Ctla2a
Dusp10
Cd2
Il18r1
Carmil2
Slamf6
Psmb10
Tbx21
Rpl22
Aif1
H2−Ab1
H2−Aa
Cd74
Nlrp3
Irf4
Cd8a
Ccl5
Spn
Lat
Batf
Lck
Gli3
Igf1
Sash3
Coro1a
Itgal
Il12rb1
Ptger4
Nfkbid
Cd3g
Laptm5
Lfng
Cd48
Dock2
Nlrc3
Ccdc88b
Rasal3
Il1b
Ptprc
Tnfrsf21
Il1rl2
Cd5
Lef1
Fcer1g
Gja1
Cd6
Gm5150
Runx2
Apbb1ip
Sirpb1c
Prdm1
Sirpb1b
Runx1
Prnp
Ada
Bcl2a1d
Zfp36l1
Gadd45g
Lmo1
Cd3e
Psap
1.5
1
0.5
0
−0.5
−1
−1.5
Garbani
No_Infection
Dm28c
TcLu
TcKr

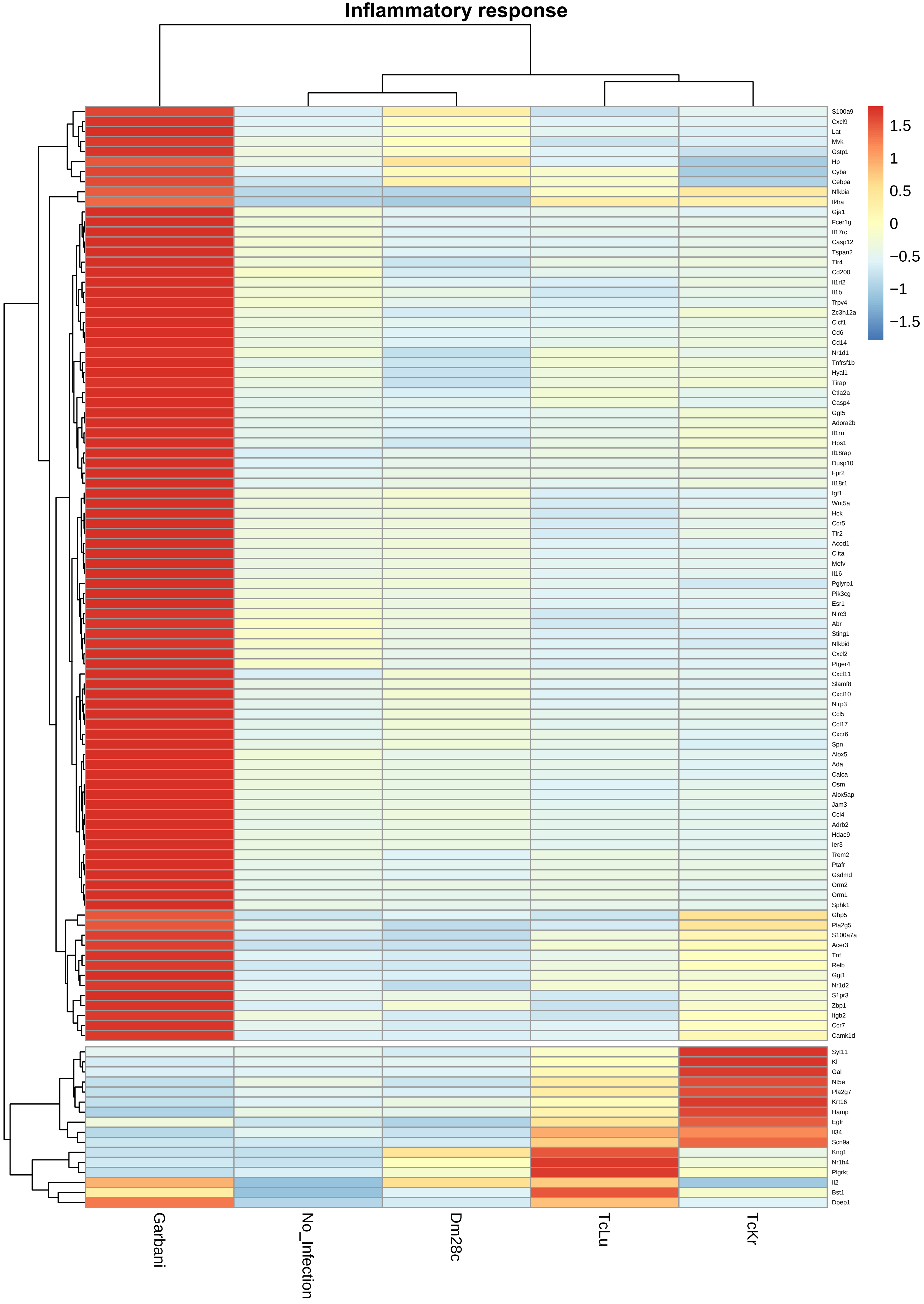
Inflammatory response
1.5
1
0.5
0
−0.5
−1
−1.5
S100a9
Cxcl9
Lat
Mvk
Gstp1
Hp
Cyba
Cebpa
Nfkbia
Il4ra
Gja1
Fcer1g
Il17rc
Casp12
Tspan2
Tlr4
Cd200
Il1rl2
Il1b
Trpv4
Zc3h12a
Clcf1
Cd6
Cd14
Nr1d1
Tnfrsf1b
Hyal1
Tirap
Ctla2a
Casp4
Ggt5
Adora2b
Il1rn
Hps1
Il18rap
Dusp10
Fpr2
Il18r1
Igf1
Wnt5a
Hck
Ccr5
Tlr2
Acod1
Ciita
Mefv
Il16
Pglyrp1
Pik3cg
Esr1
Nlrc3
Abr
Sting1
Nfkbid
Cxcl2
Ptger4
Cxcl11
Slamf8
Cxcl10
Nlrp3
Ccl5
Ccl17
Cxcr6
Spn
Alox5
Ada
Calca
Osm
Alox5ap
Jam3
Ccl4
Adrb2
Hdac9
Ier3
Trem2
Ptafr
Gsdmd
Orm2
Orm1
Sphk1
Gbp5
Pla2g5
S100a7a
Acer3
Tnf
Relb
Ggt1
Nr1d2
S1pr3
Zbp1
Itgb2
Ccr7
Camk1d
Syt11
Kl
Gal
Nt5e
Pla2g7
Krt16
Hamp
Egfr
Il34
Scn9a
Kng1
Nr1h4
Plgrkt
Il2
Bst1
Dpep1
Garbani
No_Infection
Dm28c
TcLu
TcKr

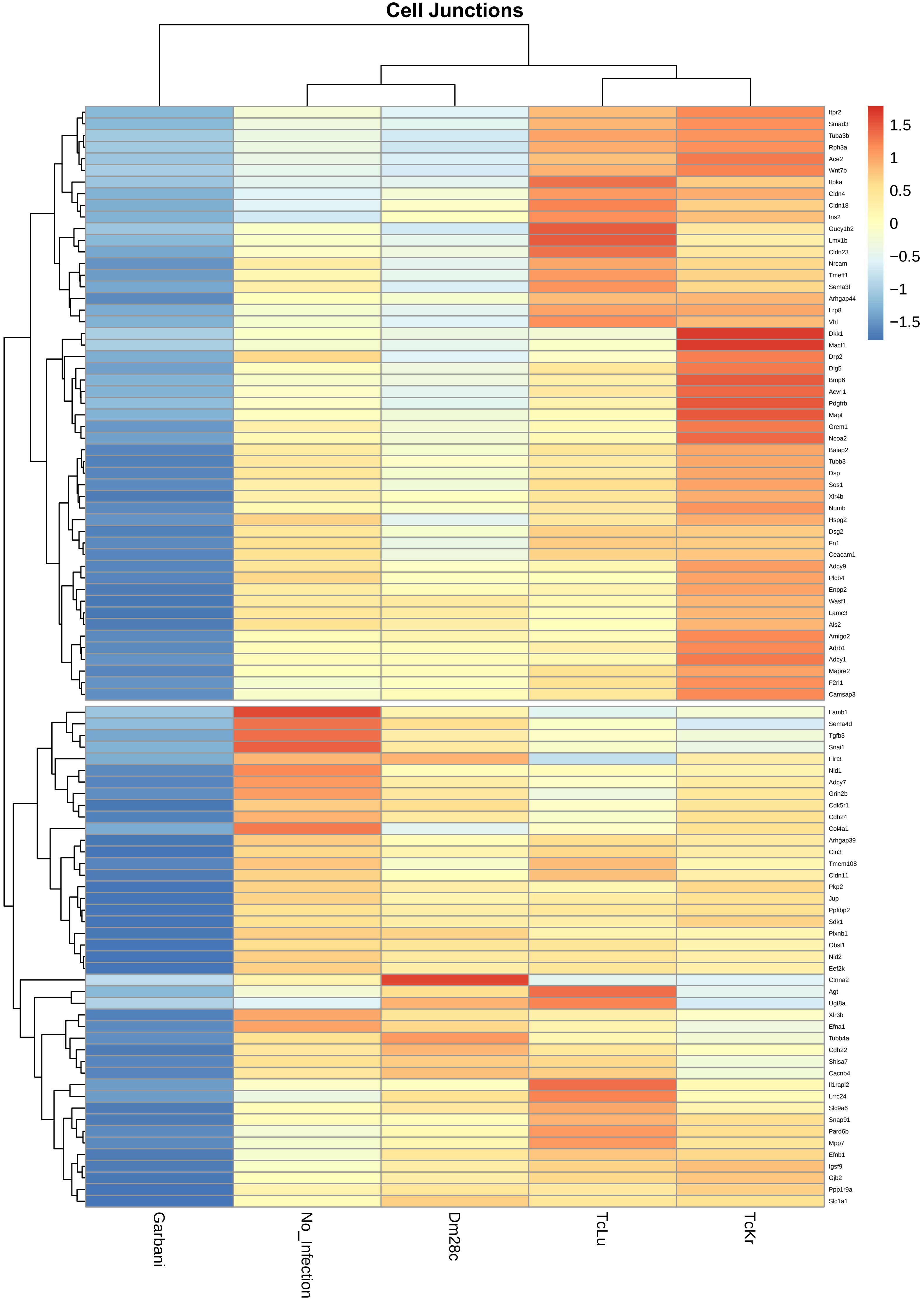

Cell Junctions
Itpr2
Smad3
Tuba3b
Rph3a
Ace2
Wnt7b
Itpka
Cldn4
Cldn18
Ins2
Gucy1b2
Lmx1b
Cldn23
Nrcam
Tmeff1
Sema3f
Arhgap44
Lrp8
Vhl
Dkk1
Macf1
Drp2
Dlg5
Bmp6
Acvrl1
Pdgfrb
Mapt
Grem1
Ncoa2
Baiap2
Tubb3
Dsp
Sos1
Xlr4b
Numb
Hspg2
Dsg2
Fn1
Ceacam1
Adcy9
Plcb4
Enpp2
Wasf1
Lamc3
Als2
Amigo2
Adrb1
Adcy1
Mapre2
F2rl1
Camsap3
Lamb1
Sema4d
Tgfb3
Snai1
Flrt3
Nid1
Adcy7
Grin2b
Cdk5r1
Cdh24
Col4a1
Arhgap39
Cln3
Tmem108
Cldn11
Pkp2
Jup
Ppfibp2
Sdk1
Plxnb1
Obsl1
Nid2
Eef2k
Ctnna2
Agt
Ugt8a
Xlr3b
Efna1
Tubb4a
Cdh22
Shisa7
Cacnb4
Il1rapl2
Lrrc24
Slc9a6
Snap91
Pard6b
Mpp7
Efnb1
Igsf9
Gjb2
Ppp1r9a
Slc1a1
1.5
1
0.5
0
−0.5
−1
−1.5
Garbani
No_Infection
Dm28c
TcLu
TcKr

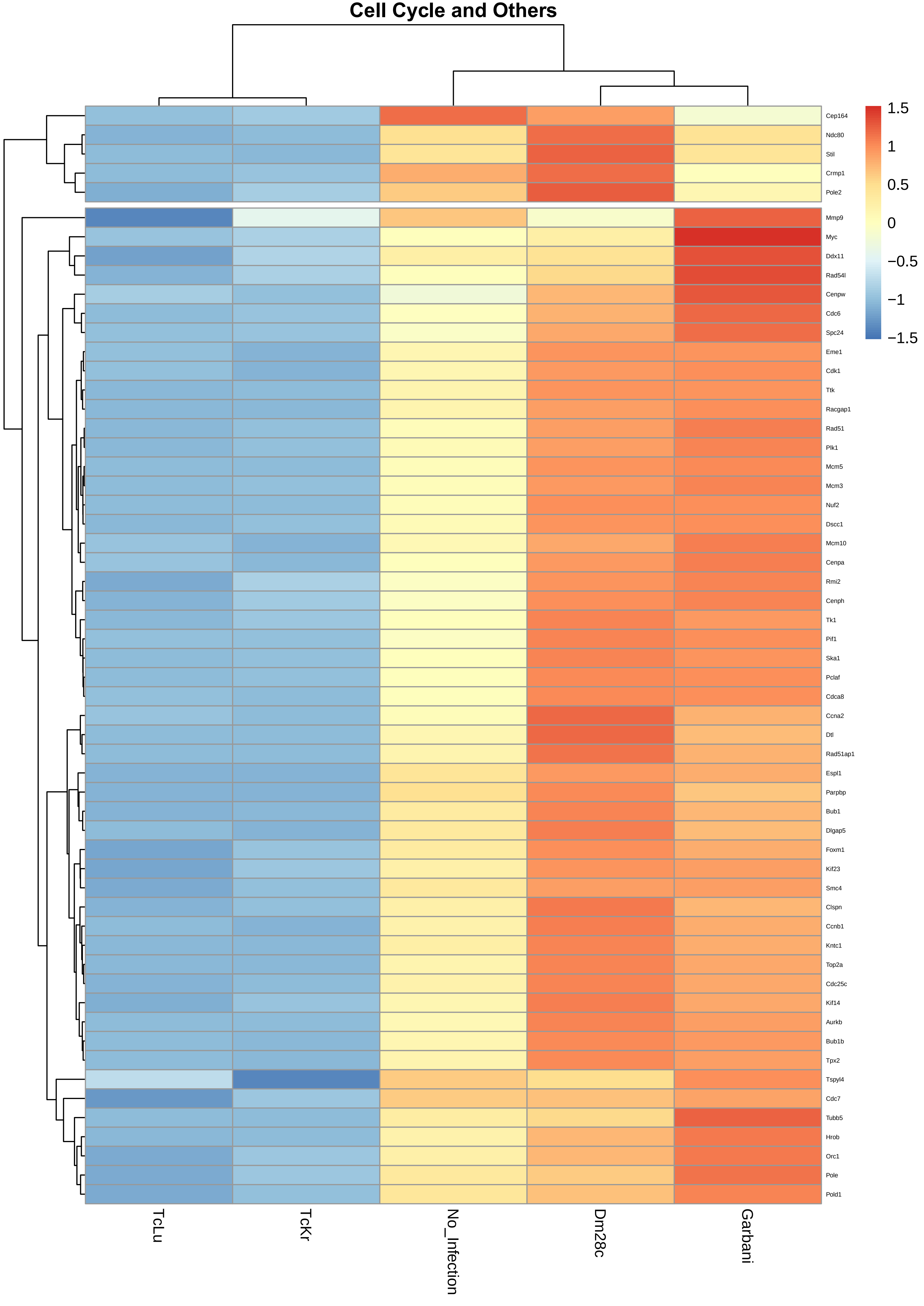

Cell Cycle and Others
1.5
1
0.5
0
−0.5
−1
−1.5
Cep164
Ndc80
Stil
Crmp1
Pole2
Mmp9
Myc
Ddx11
Rad54l
Cenpw
Cdc6
Spc24
Eme1
Cdk1
Ttk
Racgap1
Rad51
Plk1
Mcm5
Mcm3
Nuf2
Dscc1
Mcm10
Cenpa
Rmi2
Cenph
Tk1
Pif1
Ska1
Pclaf
Cdca8
Ccna2
Dtl
Rad51ap1
Espl1
Parpbp
Bub1
Dlgap5
Foxm1
Kif23
Smc4
Clspn
Ccnb1
Kntc1
Top2a
Cdc25c
Kif14
Aurkb
Bub1b
Tpx2
Tspyl4
Cdc7
Tubb5
Hrob
Orc1
Pole
Pold1
TcLu
TcKr
No_Infection
Dm28c
Garbani

**Mitotic Cell Cycle Process**

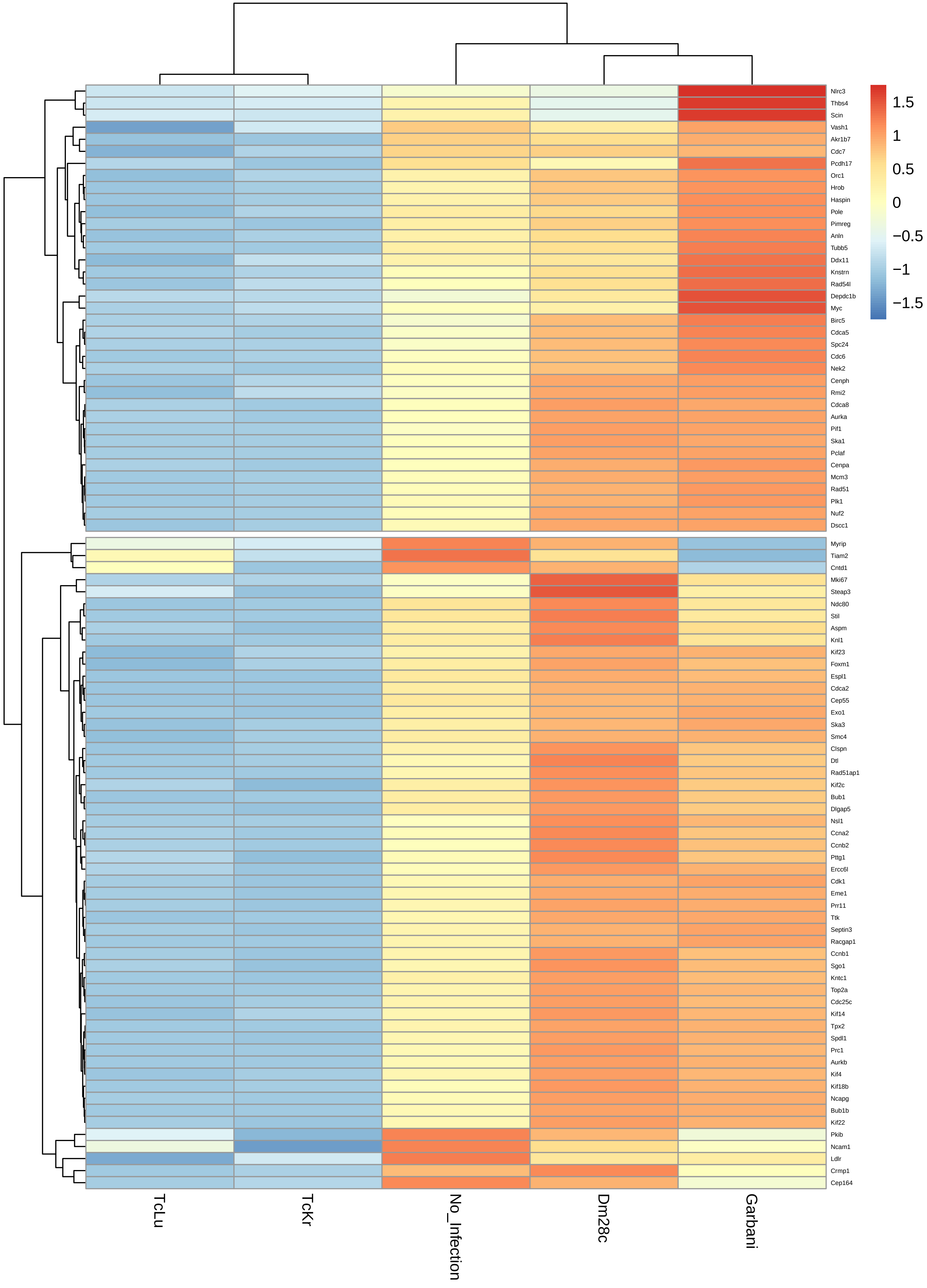

### Cell Projections and Morphogenesis

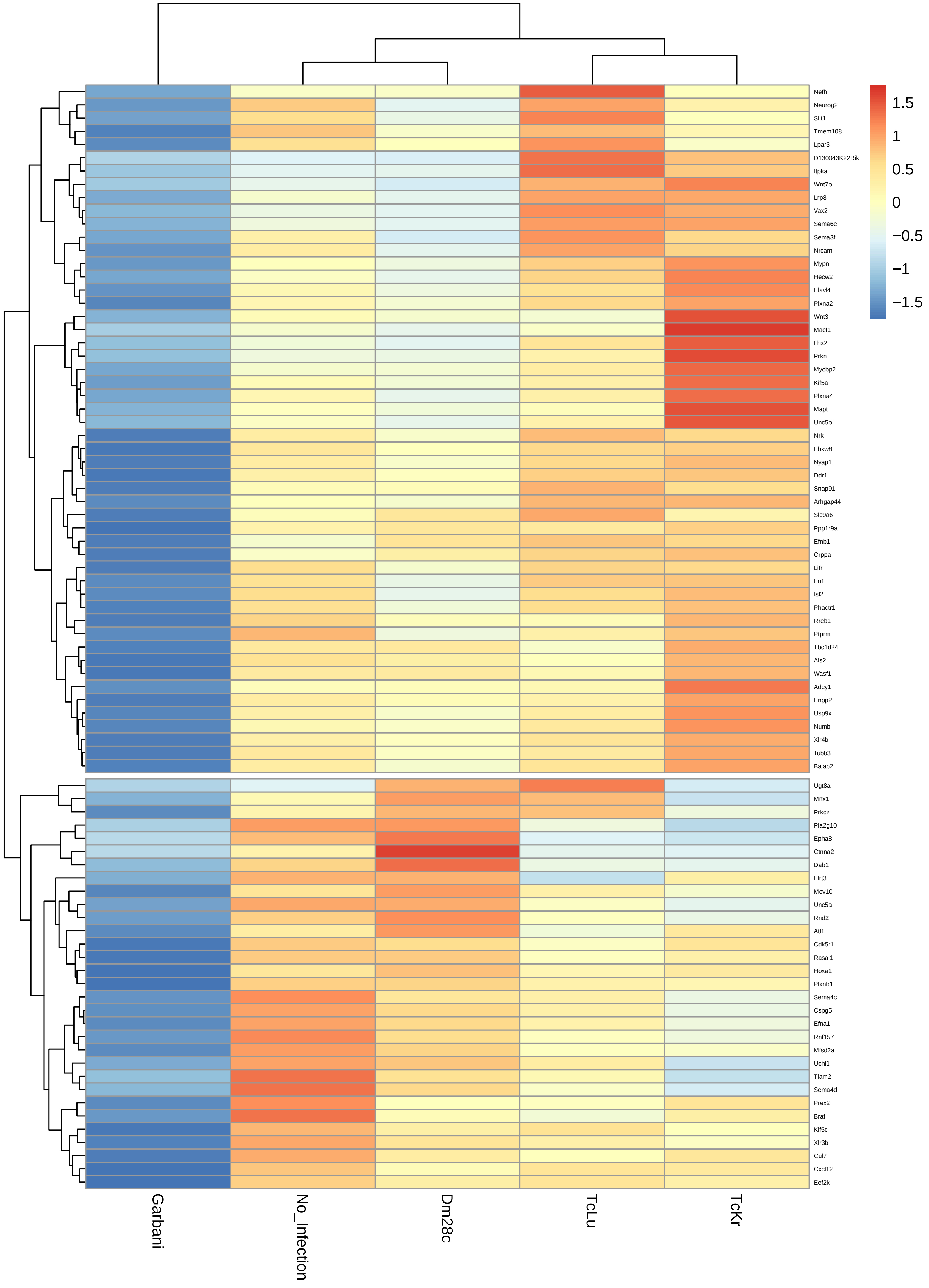

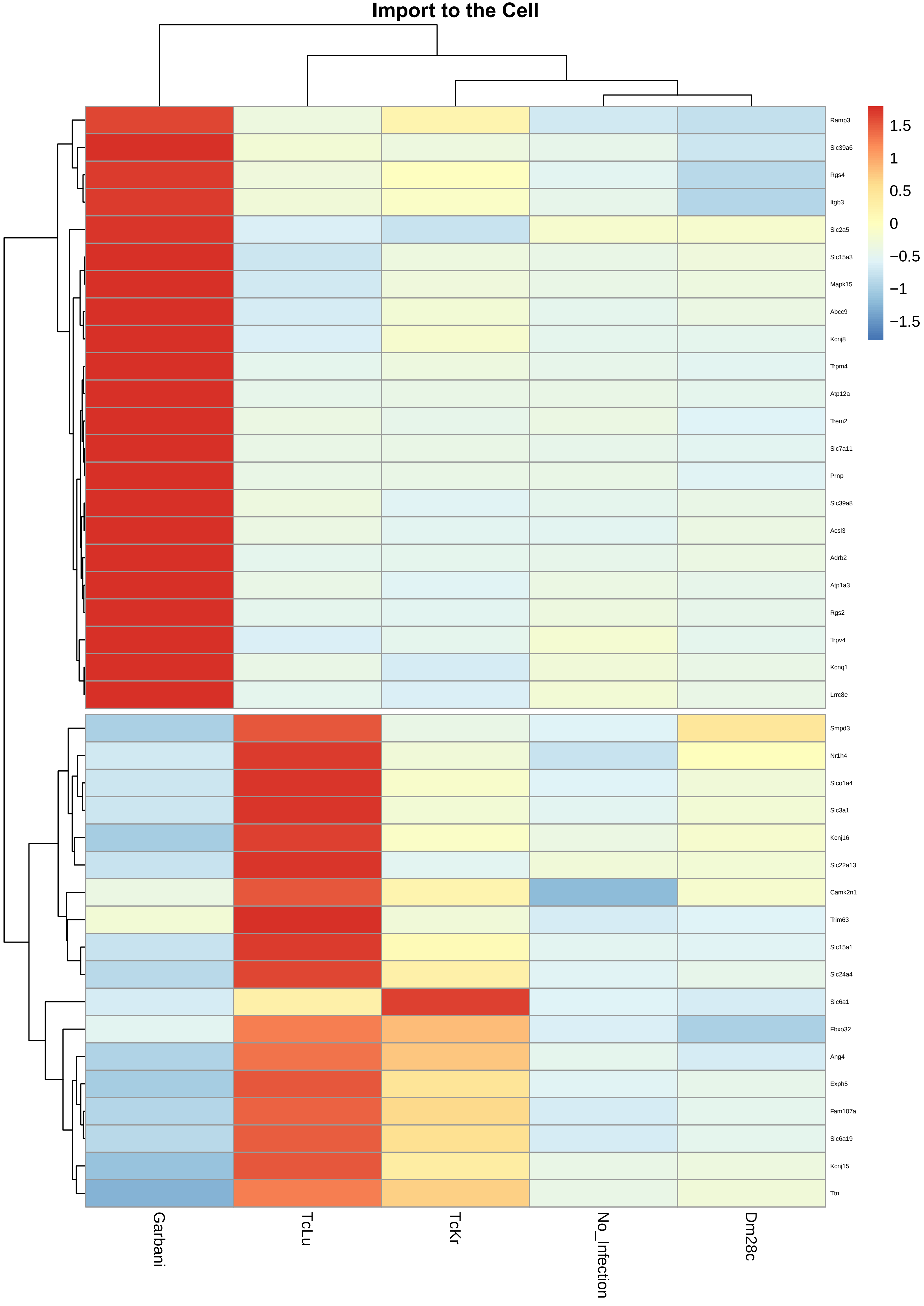
Import to the Cell
Ramp3
Slc39a6
Rgs4
Itgb3
Slc2a5
Slc15a3
Mapk15
Abcc9
Kcnj8
Trpm4
Atp12a
Trem2
Slc7a11
Prnp
Slc39a8
Acsl3
Adrb2
Atp1a3
Rgs2
Trpv4
Kcnq1
Lrrc8e
Smpd3
Nr1h4
Slco1a4
Slc3a1
Kcnj16
Slc22a13
Camk2n1
Trim63
Slc15a1
Slc24a4
Slc6a1
Fbxo32
Ang4
Exph5
Fam107a
Slc6a19
Kcnj15
Ttn
Garbani
TcLu
TcKr
No_Infection
Dm28c
1.5
1
0.5
0
−0.5
−1
−1.5

### Positive Regulation of Secretion and Import

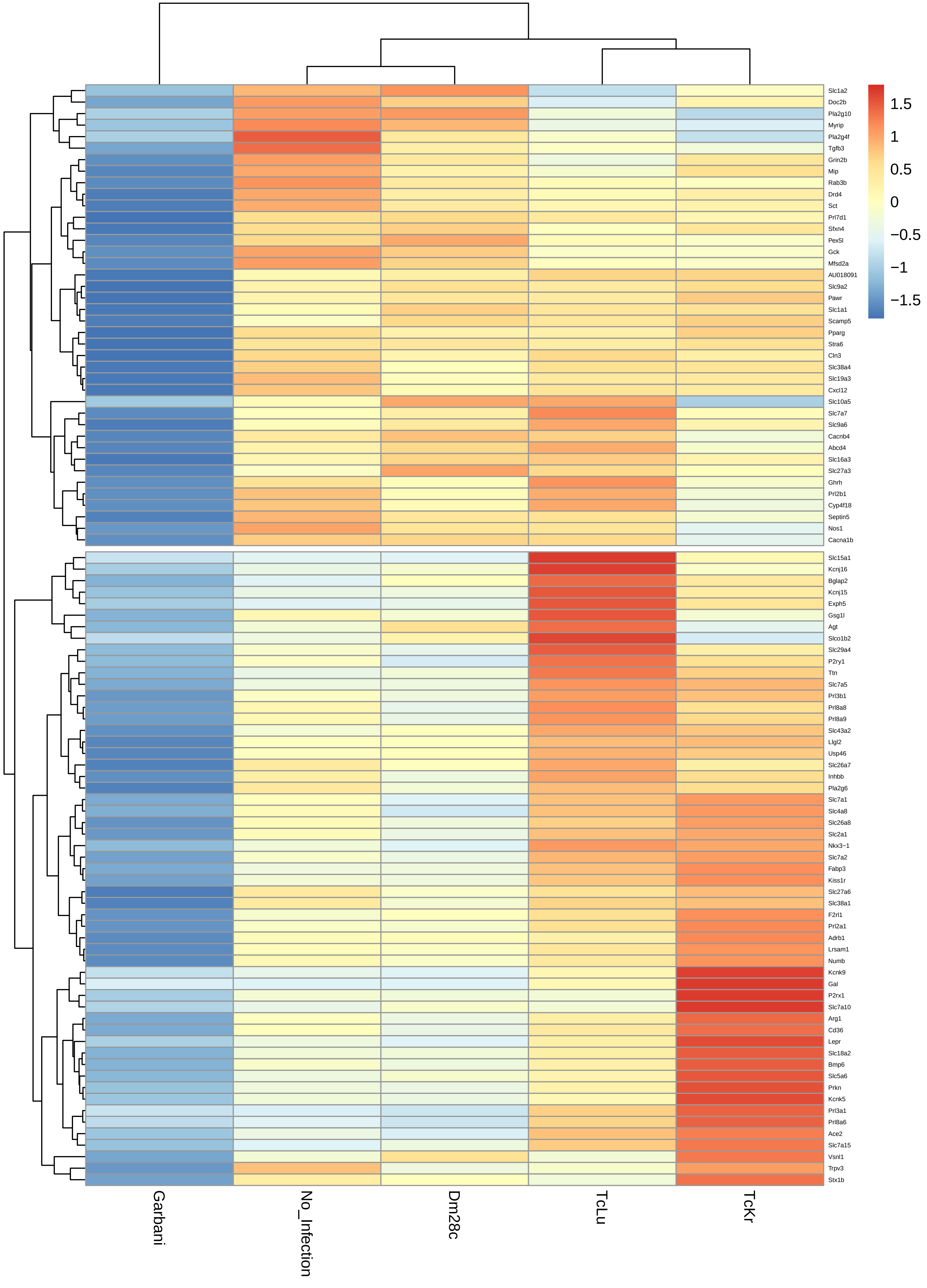

### Positive Regulation of Secretion

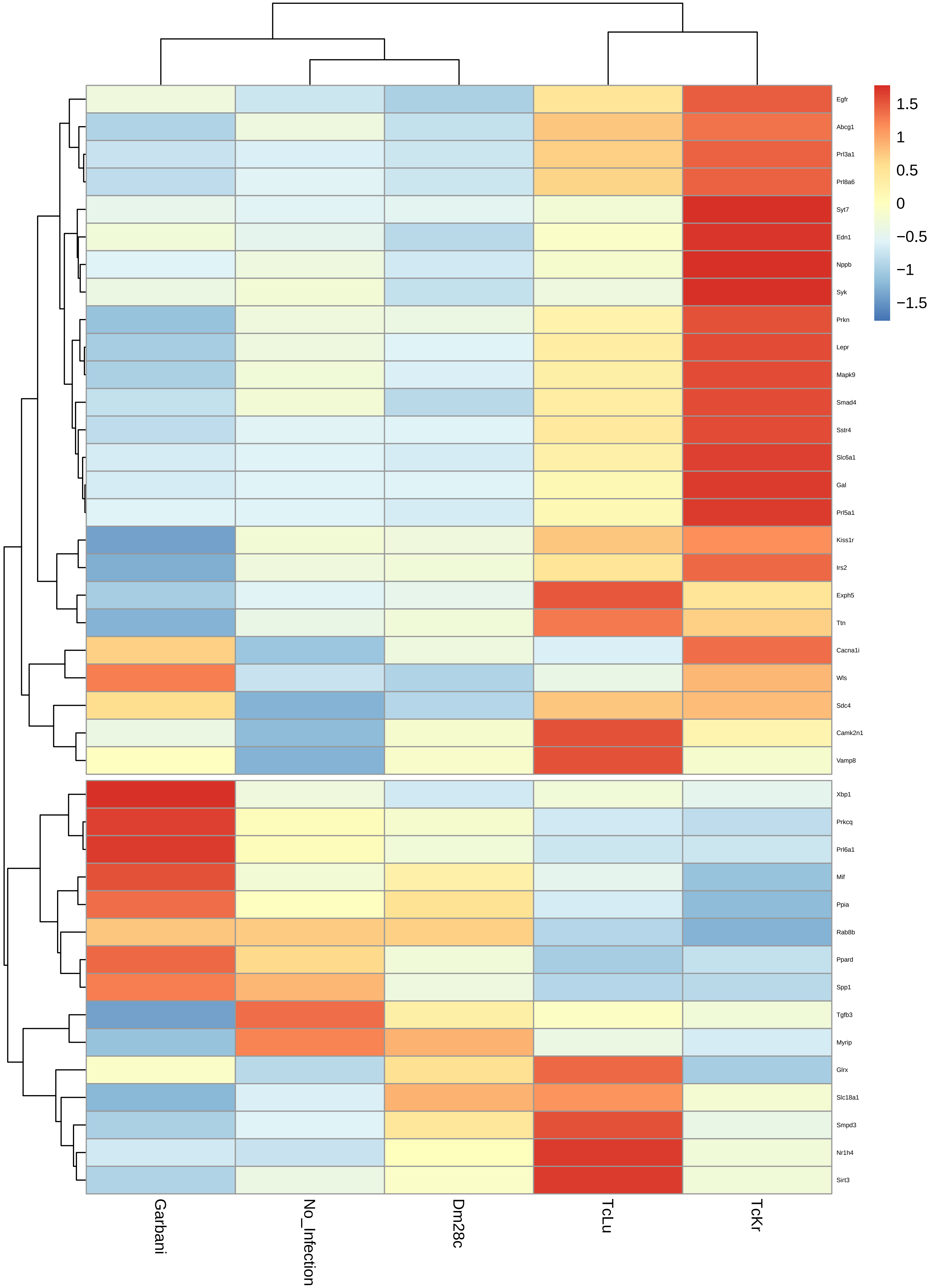

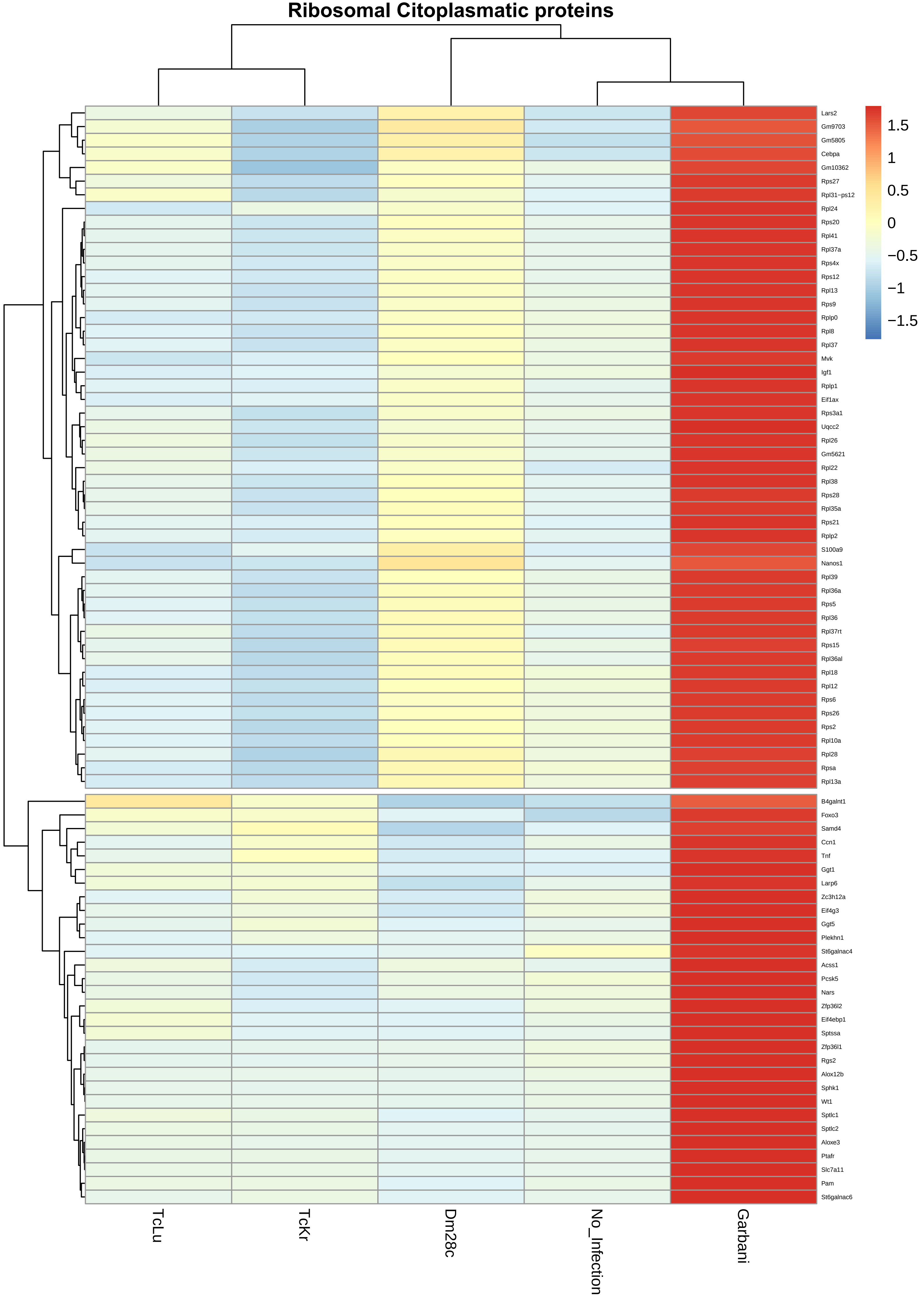
Ribosomal Citoplasmatic proteins
1.5
1
0.5
0
−0.5
−1
−1.5
Lars2
Gm9703
Gm5805
Cebpa
Gm10362
Rps27
Rpl31−ps12
Rpl24
Rps20
Rpl41
Rpl37a
Rps4x
Rps12
Rpl13
Rps9
Rplp0
Rpl8
Rpl37
Mvk
Igf1
Rplp1
Eif1ax
Rps3a1
Uqcc2
Rpl26
Gm5621
Rpl22
Rpl38
Rps28
Rpl35a
Rps21
Rplp2
S100a9
Nanos1
Rpl39
Rpl36a
Rps5
Rpl36
Rpl37rt
Rps15
Rpl36al
Rpl18
Rpl12
Rps6
Rps26
Rps2
Rpl10a
Rpl28
Rpsa
Rpl13a
B4galnt1
Foxo3
Samd4
Ccn1
Tnf
Ggt1
Larp6
Zc3h12a
Eif4g3
Ggt5
Plekhn1
St6galnac4
Acss1
Pcsk5
Nars
Zfp36l2
Eif4ebp1
Sptssa
Zfp36l1
Rgs2
Alox12b
Sphk1
Wt1
Sptlc1
Sptlc2
Aloxe3
Ptafr
Slc7a11
Pam
St6galnac6
TcLu
TcKr
Dm28c
No_Infection
Garbani
